# An integrated enzymatic and computational pipeline for quantifying off-target base-editing

**DOI:** 10.1101/2025.08.26.667396

**Authors:** Alexander G. McFarland, Soon-Keat Ooi, Davin Tafuri, Elahe Kamali, Natalie Cooper, Emma J. Cook, Shantan Reddy, Danuta Jarocha, Nils Wellhausen, Saar I. Gill, Aoife M. Roche, Joseph A. Fraietta, Frederic D. Bushman, Friederike Herbst

**Affiliations:** Department of Microbiology, Perelman School of Medicine, University of Pennsylvania, Philadelphia, PA 19104; Center for Cellular Immunotherapies, Perelman School of Medicine, University of Pennsylvania, Philadelphia, PA 19104; Division of Hematology/Oncology, Department of Medicine, University of Pennsylvania, Philadelphia, PA 19104; Department of Pathology and Laboratory Medicine, University of Pennsylvania, Philadelphia, PA 19104

**Keywords:** DNA base editing, CRISPR/Cas9, gene modification, ABE8e, off-target analysis, iGUIDE, cell-type specific off-target sites

## Abstract

DNA base editing is increasingly used for human genetic modification, but methods for monitoring off-target editing are nascent. Here we present a simple model-independent workflow for identifying sites of off-target base-editing in relevant cell types on a genome-wide level. We report that sites of off-target editing by the ABE8e editor could be identified using an ABE8e derivative with restored DSB cleavage activity. This allows marking of enzyme-generated double-stranded (ds) DNA breaks by incorporation of dsDNA oligonucleotides that are transfected into primary target cells. DNA sequencing at sites of oligonucleotide incorporation reported both the genomic location of off-target cleavage and the extent of base-editing nearby. We present a platform combining this cellular (BEiGUIDE-Seq) and computational workflow (CRISPRito) to generate optimized amplicon panels for convenient monitoring of off-target base editing. This work introduces a generalizable strategy to evaluate off-target edits in patient-derived cells, addressing a critical safety gap for clinical base editing.

## Introduction

Methods are now well developed for editing pre-specified bases in the human genome. One widely used method involves fusion of a CRISPR/Cas9 RNA-guided binding unit to a DNA deaminase, for example APOBEC ^1^ or TadA ^2,3^. In these enzymes, the Cas9 protein is modified to reduce the normal double-strand DNA cleavage activity, but single-stranded DNA cleavage is preserved to specify the template strand chosen for DNA repair after editing. DNA base editors have been extensively optimized for activity and specificity, as with the adenine base editor ABE8e ^2–10^. Base editing strategies have been developed to treat sickle cell disease ^11^, beta thalassemia, hypercholesterolemia ^12–14^, T cell leukemia ^15^, and rare diseases such as severe carbamoyl-phosphate synthetase 1 deficiency ^2, 16^.

Here we present a generalized method for monitoring off-target editing for use in preclinical and clinical applications. As a model, we used the adenine base editor ABE8e and targeted the *PTPRC* gene. Genetic engineering of the chimeric antigen receptor (CAR) binding epitope in CD45 (encoded by *PTPRC*) prevents engineered CAR-T cells from attacking each other (“fratricide”) and leukopenia by shielding benign hematopoietic blood cells through epitope editing of CD34+ cells ^17^.

Multiple methods have been developed for monitoring off-target editing (Supplementary Table 1). Many rely on in silico nomination or biochemical assays using naked DNA; such methods do not assess outcomes in the target cell type and do not take into account genetic polymorphism in the specific patient to be treated.

Here sites of potential off-target activity are nominated using a modified version of ABE8e in which the SpCas9 dsDNA cleavage capacity has been restored (termed ABE8e-Cas9^WT^), allowing marking of sites of enzyme binding and dsDNA cleavage by oligonucleotide incorporation (iGUIDE method) ^18^. Sequencing sites of oligonucleotide incorporation also reports base editing nearby. This allows convenient interrogation of off-target sites in pertinent cell types, thereby documenting possible effects of chromatin architecture and human polymorphism (Fig. 1A). Added to these nominated sites are sites proposed from a variety of bioinformatic tools, with the highest priority given to the most highly ranked and those in locations that could potentially result in genotoxicity. We present a deep analysis of 94 samples of ex vivo modified cells from 12 human donors, and software packages (Fig. 1B) for analyzing editing at sites of oligonucleotide incorporation (BEiGUIDE) and for combining data sets to provide a prioritized list of candidate off-target sites for convenient analysis by amplicon sequencing (CRISPRito).

**Figure 1.**
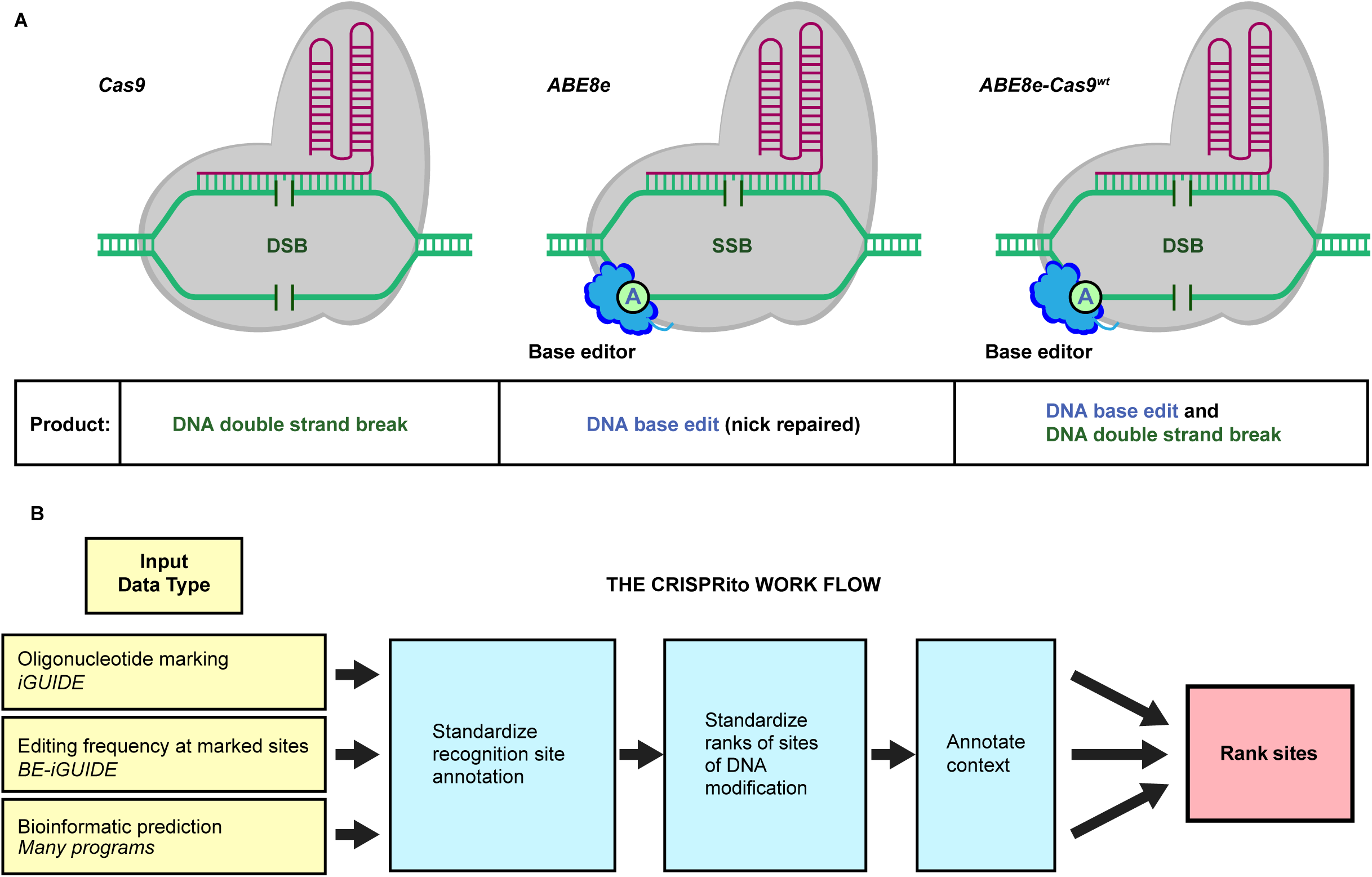
Diagram of the Cas9 derivatives studied, and the CRISPRito analytical pipeline. A) Activities of three enzymes were compared. These are i) wild-type Cas9, ii) ABE8e, which is comprised of a base editor (TadA domain, BE) and the Cas9 binding unit, with one DNA nuclease domain inactivated by mutation (Cas9^mut^), and iii) ABE8e-Cas9^WT^, which is the same as the ABE8e editor, except both nuclease domains in the Cas9 domain are active. B) The CRISPRito pipeline. Starting from the left, several forms of data are introduced into the pipeline: ABE8e-Cas9^WT^ oligonucleotide marking analyzed by iGUIDE data, editing data, and data from bioinformatic predictions. The sgRNA recognition sites are standardized, and sites are ranked as called by each method. The context of each site is also annotated (predictions of editing within an exon, or editing in or near a cancer-associated gene). All the data is then used to create a single ranked list for generation of an amplicon panel for analysis of specified sites.

## Results

### Analyzing sites of off-target activity of ABE8e and derivatives

To assess the on-target and genome-wide off-target sites of editing by ABE8e and derivatives in vivo, we first assessed sites of DNA double-strand cleavage by marking dsDNA breaks with incorporated oligonucleotides (Fig. 2). For this, we used the iGUIDE pipeline ^18^, a modification of GUIDE-seq ^19^ with improved specificity. In this approach, protected double-stranded oligonucleotides (dsODN) were introduced into primary T cells or CD34+ cells together with the gene modifying machinery, which was composed of the *PTPRC* gRNA with one of three enzymes (Fig. 1A): the standard ABE8e nickase; a re-engineered version with restored dsDNA cleavage activity--named ABE8e-Cas9^WT^--to allow for simultaneous base editing and efficient DNA double-strand break generation; or wild type Cas9. DNA was then isolated from gene-modified and dsODN-tagged cells. Genomic DNA molecules were sheared, and DNA adaptors were ligated onto the free DNA ends, allowing PCR amplification of the dsODN-marked regions. PCR products were then sequenced. Typically, ABE8e supports only single-stranded DNA cleavage due to a mutation in the RuvC-like Cas9 domain ^20^ leading to low DSB activity ^4, 21, 22^. However, we observed low levels of DNA oligonucleotide incorporation with ABE8e, indicating rare DNA breakage on the other strand near the nick. We compared iGUIDE results using ABE8e, ABE8e-Cas9^WT^, and conventional Cas9. We tested two different time points for DNA harvest and three concentrations of the dsODN (details in Supplementary Tables 2 and 3). We found little difference, and so treated each data set as a separate experimental replicate. In a few cases, a second guide was compared. The cells analyzed were obtained from 12 normal human donors (Supplementary Table 4).

**Figure 2.**
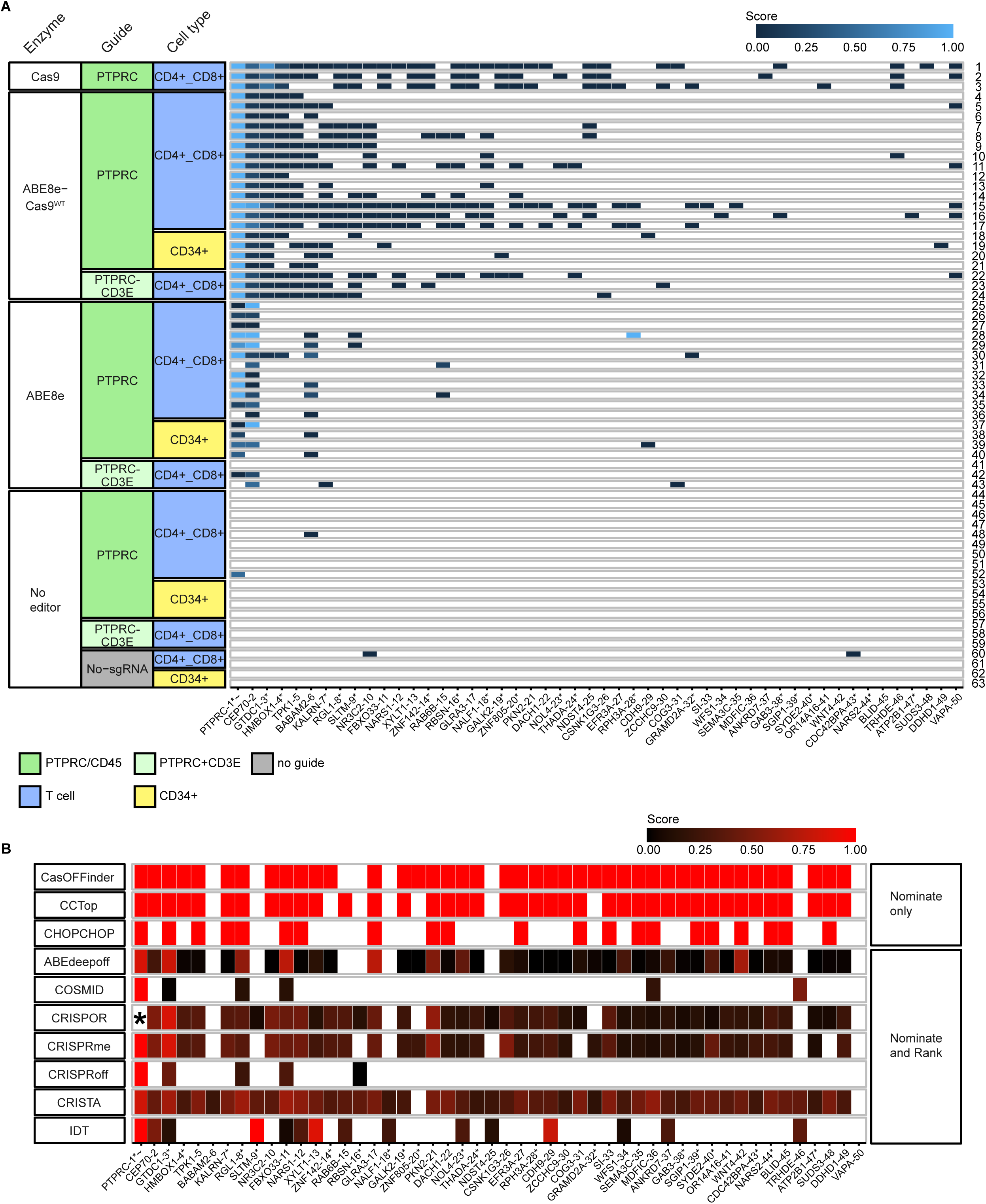
On-target and off-target activity queried using oligonucleotide incorporation at sites of dsDNA breaks (iGUIDE). A) Summary of marking by the iGUIDE method at on-target and off-target sites. Each row shows results for an individual sample, each column in the heat map shows the efficiency of marking for a different target site. Each cell is scored by the efficiency of marking (key at bottom). Each site is named by the nearest gene (row along bottom). Asterisks and tildes indicate the site is in a transcription unit and in an oncogene, respectively. The enzymes used were (from the top) Cas9, ABE8e-Cas9^WT^, ABE8e, and no editor. The guides used targeted *PTPRC* (CD45), or both *PTPRC* and *CD3E*. Cell types were either T cells (CD4+ CD8+) or hematopoietic stem cells (CD34+). B) Comparison of off-target sites called by each of 10 bioinformatic tools. Each row shows results for a different tool, each column shows scores for different sites, named by the nearest gene. The tools called an average of 647 off-target sites (range 11-3,181). The top three tools did not provide relative rankings, so all sites are called equally.

Figure 2 shows examples of locations detected for dsODN incorporation in cells treated with Cas9, ABE8e-Cas9^WT^, or ABE8e (Fig. 2A). Sites of dsODN incorporation were cataloged for each of the 63 samples tested. The relative use of each site was quantified by comparing the number of independent adaptor positions detected in reads for each different chromosomal target. Adaptors mark each DNA chain with a unique position because the sonication step results in DSBs at random sites. This allows quantification of the relative numbers of cells sampled using the number of independent adaptor positions associated with each dsODN position.

For ABE8e-Cas9^WT^ and Cas9, the on-target *PTPRC* site showed the highest frequencies of oligonucleotide incorporation in every sample (n=24; 18.1%-89.3 %). No differences were seen between ABE8e-Cas9^WT^ and Cas9 on-target frequencies in a comparison over three donors (Supplementary Fig. 1). Each of these samples also reported the same two top off-target sites, near the gene *CEP70*, and within an intron of *GTDC1*. Frequencies of the top five off-target sites, which were shared by all seven donors in the ABE8e-Cas9^WT^data, ranged from 2.1±1.7% to 36.7±27.0%, normalized to the *PTPRC* on-target site detection (Supplementary Table 5). An additional 14 potential off-target sites, which were detected in at least three out of seven donors, were all below 1% relative to *PTPRC* (Supplementary Table 6). A wide range of minor sites was also detected in individual samples. No obvious differences were seen for editing at these sites when a second guide RNA was added. Overall, a total of eight sites were detected in at least half of the pooled ABE8e-Cas9^WT^ plus Cas9 samples (Fig. 2A).

Data was much sparser for the ABE8e editor; here, many samples identified the top *PTPRC* site, and the most frequent off-target site, *CEP70*. In the no-editor controls, we only recovered sites marked by single oligonucleotide incorporations, likely corresponding to integration at naturally occurring dsDNA breaks.

### Comparison of bioinformatic methods for identifying sites of off-target cleavage and base editing

We next compared predicted off-target locations using ten bioinformatic tools designed to predict sites of Cas9 cleavage (Fig. 2B and Supplementary Table 7). All methods identified the *PTPRC* site as a top candidate. The bioinformatic tools called a large number of additional candidate sites, from 11 (COSMID) to 3,183 (CRISTA). These sites showed incomplete overlap with the sites reported by iGUIDE analysis, and the nominations drastically varied between the tools: only two off-target sites were predicted by all ten tools, and only 45 sites were predicted by five out of ten tools.

A principal coordinate analysis showed that results for the experimental ABE8e-Cas9^WT^ and Cas9 samples clustered tightly (Supplementary Fig. 2). The data for the unmodified ABE8e samples were scattered, indicating a low signal and contributions of oligonucleotide incorporation at spontaneous dsDNA breaks. The purely bioinformatic data sets were also widely scattered, reflecting the very large numbers of divergent calls made by each.

### Detection of base editing by the ABE8e-Cas9^WT^ enzyme at sites of oligonucleotide incorporation

The sequence data used to capture the dsODN positions could also be interrogated for frequencies of base editing (Fig. 3A). For this, a software tool (BEiGUIDE) was developed. In all samples treated with the ABE8e-Cas9^WT^ editor, editing was detectable at the *PTPRC* site, ranging from 9.6% to 61.4% of oligonucleotide-marked alleles. *CEP70* and *GTDC1* were the next most frequently edited locations, showing median editing of 55.7% and 21.3%. A few additional editing sites were detected in a minority of samples. No editing was seen at sites of dsODN incorporation in the no-editor control. For the ABE8e nickase, the recovery of sites was so sparse that no site showed editing above the 5% threshold. In addition, no editing was detected in oligonucleotide-marked sites made by conventional Cas9. Thus, editing efficiency can be quantified in the ABE8e-Cas9^WT^ iGUIDE sequence data.

**Figure 3.**
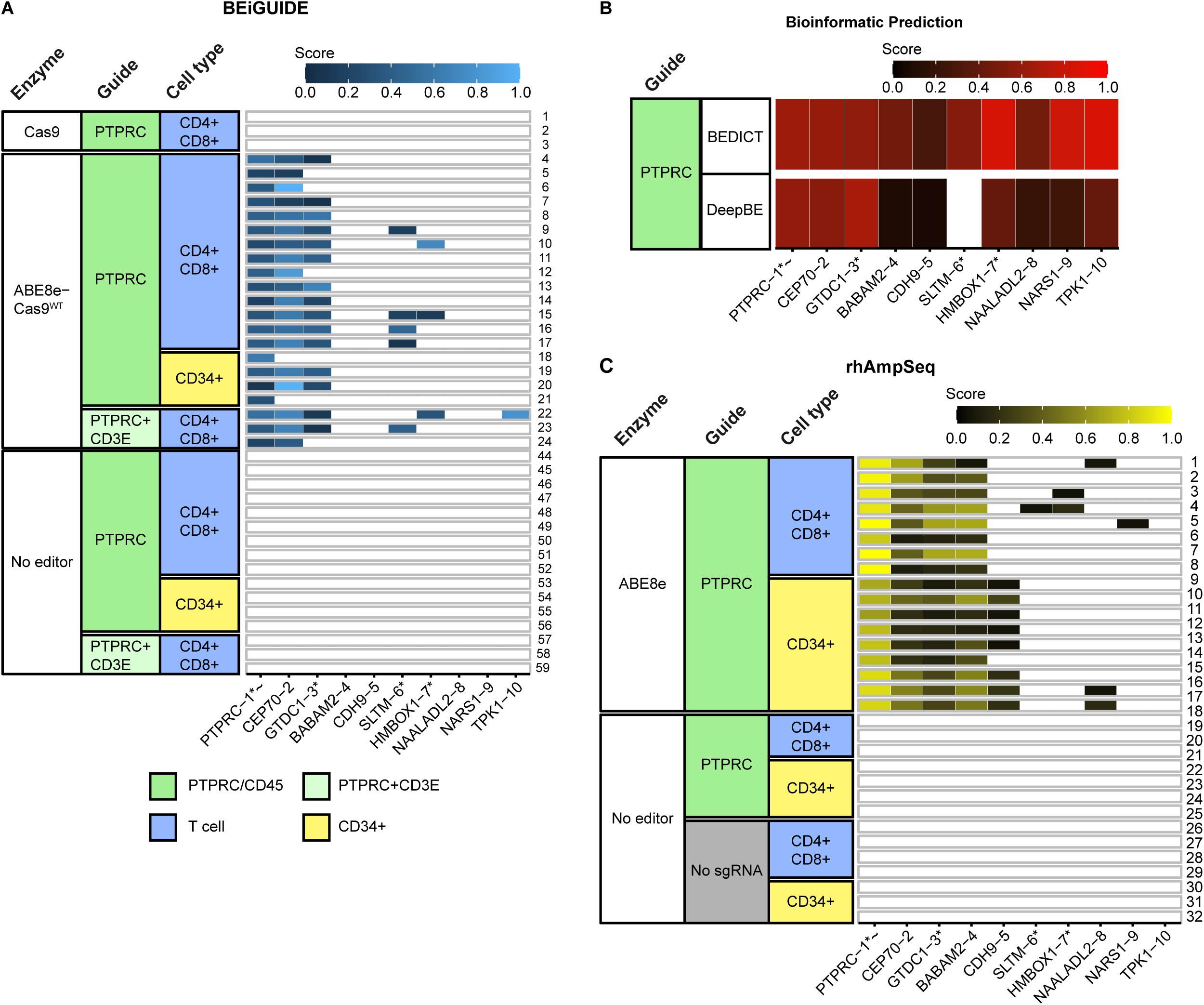
On- and off-target base editing by Cas9, ABE8e-Cas9^WT^, and ABE8e. A) On- and off-target editing measured at genomic sites marked by oligonucleotide incorporation (iGUIDE) by Cas9, ABE8e-Cas9^WT^, or no editor. The rows show different DNA specimens analyzed, the columns in the heat map show the measured extent of editing. Only sites where one base was edited in at least 5% of reads were scored; the relative extent of editing is shown by the color code (key at bottom). The enzyme, guide and cell types are color coded in the columns at the left (key at bottom). Edit sites were identified and quantified using BEiGUIDE software. B) Annotating sites of predicted base editing using bioinformatic tools. A total of six and 141 sites were called by the two tools; only those sites analyzed in part B) are shown in the figure. The relative scores are shown by the red color coding. C) On- and off-target base editing analyzed using an amplicon panel to query specific genomic loci. The amplicons to be queried were selected using CRISPRito software based on data from iGUIDE analysis of sites of oligonucleotide incorporation using ABE8e-Cas9^WT^ and Cas9; base editing by ABE8e-Cas9^WT^; and the bioinformatic tools. The extent of editing is indicated by the black to yellow coloration; other annotation as in part A). Asterisks and tildes indicate the site is in a transcription unit and in an oncogene, respectively.

### Quantifying editing efficiencies using an amplicon panel, merging information from multiple data types

We next sought to test how well the data from the above assays together could predict sites of actual base editing by the ABE8e editor. We filtered and ranked a total of 7,756 cell-based or in silico-nominated sites based on their high abundance in the iGUIDE approach and predictions by multiple in silico tools. Data was incorporated for 10 bioinformatic tools calling sites of dsDNA cleavage, and two bioinformatic tools predicting sites of base editing (Fig. 3B). The design first accepted DNA sites with empirical evidence for modification (n=87). Only 14 of these target sites overlapped with one or more bioinformatic tools. Sites were then added from those nominated by six to ten bioinformatic tools (n=69). Due to primer incompatibilities in a multiplex approach, 12 low priority sites had to be excluded, so that a total of n=144 sites were selected for further off-target validation (Supplementary Table 8).

The genomic context of edit sites is also an important consideration for patient safety, so we annotated sites in exons and cancer-associated genes. Only two out of 141 uniquely mappable sites were located in exonic regions, and these were in 3’ or 5’ UTRs. Thus, none, when edited, altered the encoded protein sequence. All others were in intronic or intergenic regions.

### Comparison of editing efficiency across the amplicon panel

We generated amplicon libraries and carried out next-generation sequencing of 31 samples, including samples from four different T cell donors; one of the T cell donors was also sampled for CD34+ cells. Fig. 3C shows the editing efficiency for ABE8e and controls. For inclusion in the figure, the sites queried needed to show at least 5% editing.

The on-target *PTPRC* site was the most frequently edited in all 17 ABE8e samples. Frequencies of editing ranged from 64.7 to 99.7% of alleles. All 17 samples reported the same three off-target sites, *CEP70*, *GTDC1*, and *BABAM2*. Editing of *CDH9* was seen selectively in 8/9 CD34+ cell samples but not in T cell samples (discussed below). A few other sites were detected in a minority of samples. All other sites queried were negative for editing at the threshold applied. The no editor controls were universally negative.

Comparison with the iGUIDE data for ABE8e-Cas9^WT^ and Cas9 showed that all four major edited off-target sites were identified by dsODN incorporation. For ABE8e-Cas9^WT^, base editing was detected at the on-target *PTPRC* site, and off-target sites *CEP70* and *GTDC1*. *BABAM2* data was less consistent (discussed below). Other sites detected by oligonucleotide incorporation (Fig. 2A), even some detected relatively robustly, were not positive in the amplicon sequence analysis (Fig. 2C). Thus, dsODN incorporation reported the sites of maximal editing identified in the amplicon panel, while most of the 144 amplicons tested were negative. A detailed analysis of precision and recall for each nomination method is presented as Supplementary Fig. 4.

The amplicon sequence data also reports the extent of editing of individual bases at each target site (Fig. 4 and Supplementary Fig. 2A-I). Editing with ABE8e (Fig. 4A) was robust at the *PTPRC* target site, with the targeted A6 residue in the protospacer sequence the most frequently edited. Several other nearby A residues were modified detectably, especially A4 and A10. However, none of these substitutions altered the encoded amino acid. The editing data was sparser with the ABE8e-Cas9^WT^ editor, so that only editing at the on-target A6 was detected in all samples (Fig. 4B).

**Figure 4.**
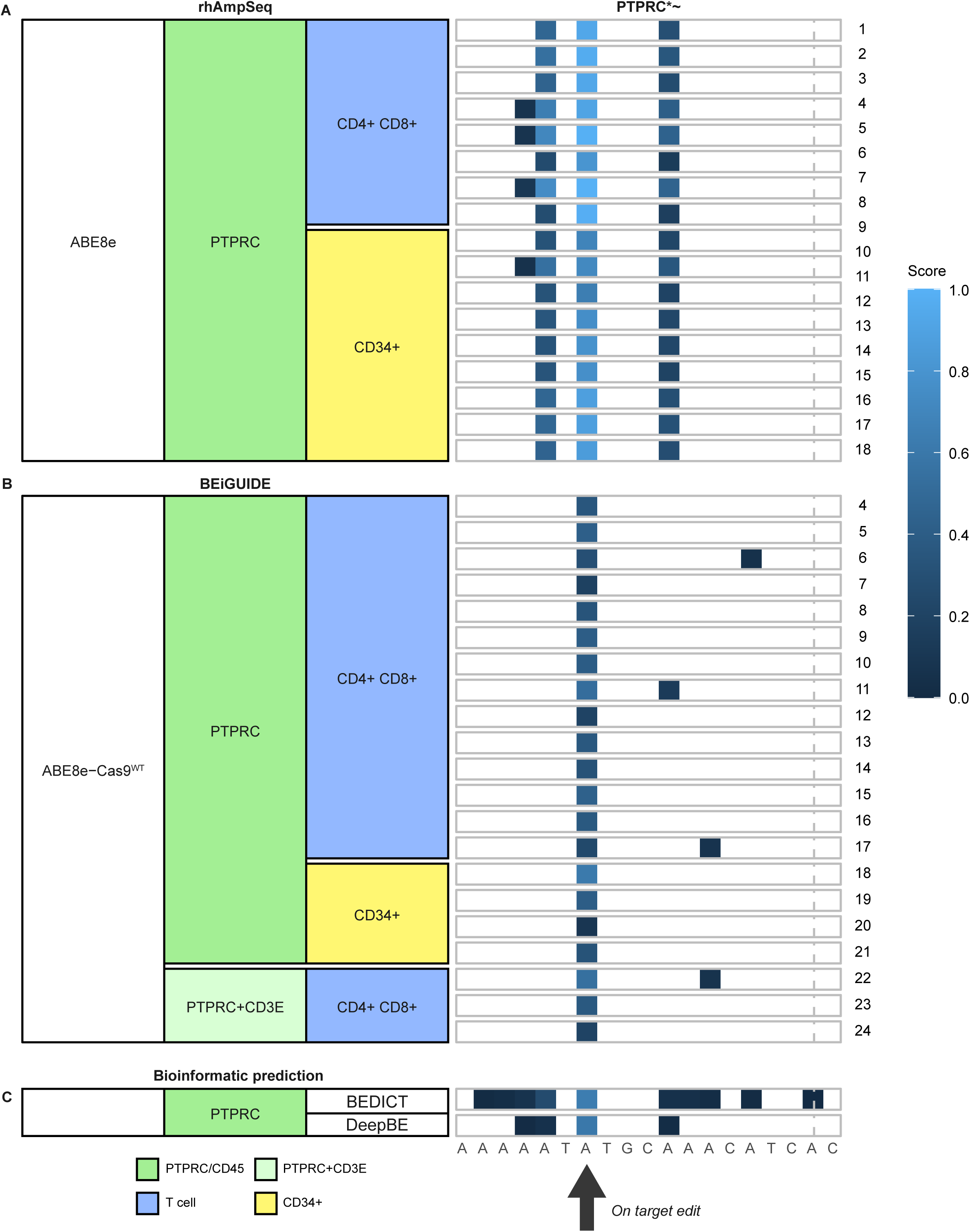
Locations of edited bases at the *PTPRC* target site. The sequence of the PTPRC target site is shown along the bottom. The black arrow shows the location of the on-target A residue. The samples analyzed are shown by the color code on the left side (columns). Samples are shown in the rows. A) Sites of base editing by the ABE8e-Cas9^WT^ at sites of oligonucleotide incorporation. B) Sites of base editing generated by the ABE8e base editing analyzed using the amplicon panel. C) Sites of base editing predicted by two bioinformatic tools. The extent of base editing is color-coded as indicated on the right. Numbering of the protospacer site starts on the left with “0”. The Cas9-targeted cut site is marked with the vertical dash. Asterisks and tildes indicate the site is in a transcription unit and in an oncogene, respectively.

Inspection of edit site distributions showed that the inconsistently marked *BABAM2* site had an unusual structure (Supplementary Fig. 4). The *BABAM2* site was heavily base-edited in the amplicon data. It was identified by only 1/10 of the bioinformatic tools querying dsDNA cleavage, but was identified by both bioinformatic tools querying base editing. It was identified sporadically in the iGUIDE oligonucleotide incorporation data. Inspection of the target site disclosed that it contains four sequential A bases, and a second run of three A bases. Thus, the site is unusually rich in targets for the base editor, possibly accounting for the differential representation in assays reporting base editing. The fact that this unusually behaved site was only identified in 1/10 of the bioinformatic methods targeting dsDNA cleavage emphasizes the value of including multiple orthogonal methods to identify off-target sites.

### Differences in editing efficiencies in different cell types from the same individual

We compared off-target editing in donor-matched T cell and CD34+ cell samples, allowing quantification of possible cell-type-specific differences. In the iGUIDE oligonucleotide marking data for Cas9 and ABE8e-Cas9^WT^ (Fig. 2A), a site in the intron of *HMBOX1* differed between the T cells (CD4+/CD8+) and CD34+ cells. Oligonucleotide incorporation was seen in all 20 T cell samples, but only in 1/4 CD34+ cell samples (Fisher’s Exact Test nominal p value of p=0.0002). For the unmodified ABE8e editor (Fig. 3B), editing at *HMBOX1* was seen in 3/17 T cell samples but in 0/4 CD34+ cells (p=nonsignificant). RNA-seq analysis of *HMBOX1* indicates that it is expressed robustly in white blood cells and lymph nodes, but not in bone marrow ^23^, indicating potential cell-type-specific accessibility.

Editing at *CDH9*, in contrast, was seen selectively in CD34+ cells (Fig. 3B). The *CDH9* site was detected in only two of the samples queried with iGUIDE (Fig. 2A), both from CD34+ cells. In contrast, *CDH9* was called by 7/10 bioinformatic tools predicting dsDNA cleavage (Fig. 2C), and by both bioinformatic tools predicting base-editing (Fig. 3C). Editing was detected in the amplicon panel assay in 8/9 CD34+ samples, but 0/8 T cell samples (nominal p=0.0004, Fisher’s exact test; Fig. 3B). RNA-seq data indicates that *CDH9* was expressed at a low level in both bone marrow and whole blood, but at higher levels in other tissues such as kidney and brain ^23^; The basis for favored editing in CD34+ cells is thus unclear. These data thus emphasize the importance of using an off-target assay querying the target cell type of interest.

No consistent differences were seen in samples from the different study participants analyzed, either in favored off-target sites or in the efficiency of base editing at the on-target site (Supplementary Fig. 1).

### CRISPRito - a bioinformatic tool to generate amplicon panels for monitoring off-target editing

To produce amplicon panels based on the above data types, we developed a software package, CRISPR data Integration TOol (CRISPRito). The package can analyze and integrate data from one or more tools, regardless of method, using a simple standardized input format. To design the panel, CRISPRito. first standardizes the genomic margins of candidate sites, which typically vary among tools. The efficiency of cleavage is then ranked. Called sites are evaluated for their presence within or near features such as exons, introns, and cancer-associated genes (and any additional user-supplied features). An editable table with configurable weights for methods, samples, and features of interest is generated to allow the user to add weights to prioritize rankings for each specific application (an example is in Supplementary Tables 9 and 10).

## Discussion

Targeted DNA base editing is an attractive tool for therapeutic human gene modification, but with implementation comes the need for methods to monitor safety. In many applications, T cells or CD34+ hematopoietic cells are genetically modified, allowing longitudinal tracking of modified cell clones by assaying cell samples from blood. It is also important to monitor off-target editing during preclinical development to support optimization of editing methods. Here we present a simple framework for monitoring off-target editing by designing optimized amplicon panels. First, sites of editing are identified biochemically. Here we introduce the use of an editor with restored dsDNA cleavage activity (ABE8e-Cas9^WT^), so that sites of off-target editing can be marked by incorporation of a double-stranded DNA oligonucleotide. Base editing can also be quantified in the resulting sequencing data. Both data types can then be merged with additional approaches, such as bioinformatic methods, to nominate sites for analysis in targeted amplicon panels using the bioinformatic pipeline CRISPRito. Genomic locations and edit sites can be interrogated for possible effects on coding sequences or nearby cancer-associated genes, and prioritized accordingly. We designed such an amplicon panel for a *PTPRC* target, and verified editing in 10 out of the 141 amplicons evaluated. Editing frequencies trailed off sharply as the supporting data became quantitatively less strong. The approach successfully distinguished a handful of true off-target edits from a long list of candidates, demonstrating a feasible method for routine surveillance of gene-edited cells. Although only ABE8e was tested, the approach is extensible to other base editors, prime editors, and multiplex editing approaches.

The cell-type-specific chromosomal landscape can influence targeting ^26^, so it is desirable to include biochemical assays in the cell type(s) of interest. Here we show consistent differences between donor-matched T cells and CD34+ cells at the *HMBOX1* and *CDH9* loci, emphasizing the value of assays in the relevant cell types. Inclusion of biochemical assays using cells from the patient to be treated also provides the added advantage of reporting possible effects of human DNA polymorphism, which can also affect the function of Cas9-based tools (e. g.,27_)._

Recently Musunuru et al. ^16^ reported successful base editing to treat severe carbamoyl-phosphate synthetase 1 deficiency, and used an amplicon panel to monitor off-target activities analogous to that described here. They used GUIDE-seq to mark positions of Cas9 cleavage. This yields useful information, but use of the ABE8e-Cas9^WT^ enzyme described here allows quantification of base editing at the newly identified sites in addition to simple dsDNA cleavage, which also reports any effects of the editor deaminase domain on specificity. Musunuru et al. also used biochemical and computational assays interrogating naked chromosomal DNA (e.g., CIRCLE-seq and ONE-Seq), which can be efficient, but do not report the influence of the cellular chromatin landscape in target cells, which we report here can modulate editing.

This study has several limitations. We examined numerous experimental parameters, but only one editor. For practical reasons we limited the size of our amplicon panel to 144 amplicons, so it is possible (though unlikely) that genomic sites that were not queried might be edited. The optimal test of our method would be monitoring human clinical trials and demonstrating the ability to identify expansion of problematic clones, but to date no such events have been reported and so cannot be sampled.

In conclusion, human gene editing is advancing as a therapeutic modality, creating a need for practical safety monitoring methods. Here we describe a simple means of developing an amplicon panel for quantification of off-target editing, which will be valuable for improving the safety assessment of gene editing therapies prior to and post-approval.

## Methods

### T cell and CD34 cell isolation

CD4+ and CD8+ T cell fractions were obtained from the University of Pennsylvania Human Immunology Core. T cells and donor-matched CD34+ cells were isolated using CD4/CD8 CliniMACS beads (Miltenyi Biotec, 200-070-213-276-01, 200-070-215-275-01) followed by isolation using AutoMACS (Miltenyi Biotec) or CD34 microbeads (Miltenyi Biotec, 130-046-703) and the CliniMACS device (Miltenyi Biotec) from G-CSF mobilized leukapheresis of healthy donors obtained under a University Institutional Review Board-approved protocol.

### Primary T Cell culture

T cells were cultured in T cell complete media (TCM) (Supplemented CTS OpTmizer T Cell Expansion Media (Gibco, A10484-02), or in X-Vivo (TheraPeak Lonza, BP04-744Q), 5% Human AB Serum (Bioivt, HUMANABSRMP-HI-1c), 1% Glutamax (Gibco, 35050-061), 0.5% Penicillin Streptomycin (Gibco, 15140-122), 5 ng/mL IL-7 (Peprotech, AF-200-07-100UG), and 5 ng/mL IL-15 (Peprotech, AF-200-15-100UG).

### CD34 cell culture

Cells were cultured in StemSpan SFEM II (StemCell Technologies, 9655) supplemented with cytokines (Peprotech, AF-300-07-100UG, AF-300-019-100UG, AF-300-18-100UG, AF-200-06-100UG) 100ng/mL hSCF, 100ng/mL FLT3L, 50ng/mL TPO, 50ng/mL IL6, and 500nM UM729 (StemCell Technologies, 72332).

### Cell electroporation to deliver gene editing machinery and dsODN

CD4+ and CD8+ T cells were activated with Dynabeads Human T-Activator CD3/CD28 (Gibco, 11132D), using a bead:cell ratio of 2:1 for 48 hrs at 37°C, 5% CO_2_ before electroporation and de-beaded for electroporation.

T cells or CD34 cells were harvested, washed 1x in PBS, and re-suspended in P3 Primary Cell Nucleofector Solution with Supplement 1 (Lonza, PBP3-02250) or in Hyclone Maxcyte buffer (Cytiva, EPB1) according to the manufacturer’s instructions. The EPS solution was prepared using 8.1 µg of sgRNA (IDT or Synthego), 5 µg of mRNA encoding ABE8e, ABE8e-Cas9^WT^ or Cas9 mRNA (Trilink), and 2.5 µL 10 µM double-stranded oligonucleotide (dsODN) (IDT). The mRNAs was synthesized to contain 5’ and 3’ untranslated regions, a 5’ CleanCap, and N1-methyl-pseudouridinylated. The ABE8e coding sequence was taken from Addgene (Plasmid #138489). The Cas9 activity was restored in ABE8e by adding a A10D substitution (reversion to wild-type). 80 µL cell solution was combined with 20 µL EPS and electroporated with a Lonza 4D-Nucleofector Core Unit and X Unit, or GTx (Maxcyte). Electroporated cells were quickly transferred to plates of pre-warmed medium at 37°C, and electroporation cuvettes were washed 2x to maximize cell yield. Cells were recovered and incubated for an additional four days at 37°C, 5% CO_2_ before cell harvest and DNA isolation.

### Evaluation of on-target base editing activity and indel formation by PCR and Sanger sequencing using base editors and wtCas9 derivatives

Genomic DNA was extracted from CD4+CD8+ T or CD34+ cells 4 days post-electroporation using the Quick-DNA Miniprep Plus Kit (Zymo Research, D4069) following the manufacturer’s instructions. Genomic regions of interest were amplified using *PTPRC*-specific primers (Supplementary Table 11) and Phusion DNA Polymerase (New England Biolabs, M0530L). PCR amplicons were generated using the following cycling conditions: initial denaturation at 98°C for 30s, followed by 30 cycles of 55°C for 15s and 72°C for 15s, with final extension at 72°C for 5 min. The resulting locus-specific PCR products were purified using the DNA Clean & Concentrator-5 kit (Zymo Research, D4013). Samples were sent to Azenta/Genewiz for Sanger sequencing. .ab1 files were downloaded and analyzed using EditR ^28^ to quantify on-target editing, TIDE ^29^, and ICE ^30^ to evaluate the indel type and total frequency. Nested PCR was performed with dsODN forward primer, dsODN reverse primer, PCR forward primer, and Phusion DNA Polymerase for quality control of dsODN integration ^18^.

### Analysis of dsDNA cleavage using iGUIDE

Marking of sites of dsDNA breaks using iGUIDE was carried out as described ^18^. Briefly, cells were exposed to an enzyme generating dsDNA breaks and a dsDNA oligodeoxynucleotide (dsODN). Cells were grown to allow DNA repair and oligonucleotide incorporation at breaks. Genomic DNA was then harvested. DNA was sheared, and adaptor oligonucleotides were ligated onto the DNA ends. Amplification was then carried out using primer annealing to the dsODN and adaptor oligonucleotide. Sequences were analyzed using the Illumina platform.

A list of key reagents for biochemical analyses is presented in Supplementary Table 12.

### Bioinformatic methods for analysis of off-target cleavage sites

Published bioinformatic methods used are listed in Supplementary Table 1. Method IDT refers to CRISPR-Cas9 guide RNA design checker https://www.idtdna.com/site/order/designtool/index/CRISPR_SEQUENCE. All software was implemented as web tools according to the authors’ instructions. Tables of cut positions and scores (when available) were inputted into CRISPRito for cut site and score standardization and annotation prior to analysis. Z-standardized scores were scaled to a min-max value between zero and one for visualization purposes.

### BEiGUIDE

BEiGUIDE is a bioinformatic package written in R that extends the utility of the existing iGUIDE bioinformatic package ^18^. BEiGUIDE takes as input an iGUIDE output and the expected base edit (e.g. A->G or C->T). Internally, the base frequencies obtained from each sgRNA BAM file at a detected cut site protospacer are calculated. The frequency of each base is compared to the expected base and marked as on or off-target depending on whether the expected edit was detected. The BEiGUIDE software and user guide are freely available at https://github.com/agmcfarland/BEiGUIDE.

### Comparison between methods

Cut site profiles were compared between sites called by different enzymes or bioinformatic methods by calculating the Bray-Curtis dissimilarities based on min-max normalized scores using the vegdist function from the “vegan” package (v2.6-2) with method set to “bray”. Principal coordinate analysis was performed using the pcoa function from the “ape” package (v5.6-2) ^31^.

Concordance in the top ten ranked cut sites of samples was calculated using the cor function from the “stats” package (v4.2.2) with method set to “kendall”. This metric was calculated within each enzyme/bioinformatic group and all-versus-all between groups.

### Comparing error profiles for the different methods tested

Supplementary Fig. 3 compares the methods over all calls by compiling values for precision and recall. Calls from each method were compared to the “truth” as reported by sequencing samples treated with ABE8e and analyzed using the amplicon panel. Comparisons over the dsDNA cut plus dsODN integration method of marking sites (Supplementary Fig. 3A), emphasize that bioinformatic methods have poor precision due to overcalling edit sites, but do have good recall due to capturing a large fraction of all edited sites. Off-target site nomination by dsODN integration (iGUIDE assay) directed by the ABE8e-Cas9^WT^ enzyme, in contrast, shows higher precision and good recall, emphasizing the utility of these methods. For the nucleotide base editing data (Supplementary Fig. 3B), again, precision is low for computational methods, but recall is high. The BEiGUIDE base editing evaluation using the ABE8e-Cas9^WT^ enzyme has much lower overcalling, and so higher precision, but recall is lower due to missing some sites because the editing efficiency is lower for the dual active enzyme.

### Implementing the CRISPRito software package

CRISPRito is a tool-agnostic method developed to standardize CRISPR-induced cut site location, annotation, and scoring generated from multiple approaches and/or replicates. Typically, an amplicon panel is constructed by 1) incorporating all sites marked by both oligonucleotide incorporation and editing; 2) incorporating all sites marked by frequent oligonucleotide incorporation; 3) incorporating sites called by multiple bioinformatic methods; and 4) incorporating nominated sites at exons and/or cancer-associated genes. User input data consists of cut site position predictions and scores for each separate data source/replicate. Cut site locations are clustered using a greedy proximity-based algorithm. Each resulting cut cluster undergoes pairwise alignment of the user-provided sgRNA sequence to up to a 100 bp genomic region surrounding the likely cut position. Alignment parameters allow for DNA and RNA bulges and prioritize the expected PAM sequence. The Levenshtein distance of the spacer-protospacer alignment is then calculated. Raw scores from each entry in a cut cluster are Z-standardized and used to obtain the average Z-score, relative ranking, and 0-1 min-max value for each cut site. Cut clusters are then annotated using user-inputted coordinates of interest. For the example rank site, intron and exon ranges were obtained from RefSeq and oncogene ranges from COSMIC^32^). A table of ranking criteria weights, including intron/exon/oncogene and minimum Levenshtein distance, with default weights, is generated from the input and can be edited by the user prior to site ranking. For each cut cluster, the weights are summed and then ordered from largest to smallest to produce a ranked list.

The CRISPRito software and user guide are freely available at https://github.com/agmcfarland/CRISPRito.

### rhAMPSeq (IDT) panel design, library preparation, and sequencing

To quantify off-target editing frequencies, we performed rhAmpSeq multiplex amplicon sequencing on genomic DNA extracted from base-edited cells and negative control cells. rhAmpSeq primers were designed to amplify the on-target site and 144 off-target sites nominated by iGUIDE-seq ^18^ (Supplementary Table 6). Amplification was done using 40 ng genomic DNA per reaction with rhAmpSeq Library Mix 1 and forward and reverse assay primer pools (rhAmpSeq CRISPR Library Kit, IDT, 10007318) using the following cycling conditions: initial denaturation at 95°C for 10 min, followed by 14 cycles of 95°C for 15 s and 61°C for 8 min, with final extension at 99.5°C for 15 min. The resulting locus-specific PCR products were diluted 1:20, and 11 μl aliquots were subjected to indexing PCR using rhAmpSeq index primers and Library Mix 2 under the following conditions: 95°C for 3 min; 24 cycles of 95°C for 15 s, 60°C for 30 s, and 72°C for 30 s; followed by final extension at 72°C for 1 min. Indexed amplicons were pooled and purified prior to paired-end sequencing (150 bp) on the Illumina NextSeq 1000. Sequencing data were aligned to the human genome assembly hg38 using BWA. The resulting BAM files were then piped into BEiGUIDE to identify on and off-target base edits.

### Comparison of methods

All iGUIDE, BEiGUIDE, and computational methods were assessed for their ability to detect or predict base editing as identified by the rhAmpSeq panel. The “truth” rhAmpSeq base edits required a A->G base edit to be 5% or more abundant and found in at least three replicates. SNPs were identified in control rhAmpSeq samples and removed from consideration in the analysis. Custom scripts were used to calculate the precision and recall for each sample. For all 141 targets in the rhAmpSeq panel, BEiGUIDE samples and base editing computational tools were assessed for the presence or absence of an edit >=5% or 0.05, respectively, at any locus. iGUIDE and cut site prediction computational tools were assessed for any cut detected or predicted that coincided with an edit at any locus.

### Statistical analysis and visualization

All analyses used R v4.2.2. T-tests and Fisher’s exact tests were performed using the wilcox.test and fisher.test functions from the “stats” package (v4.2.2). Visualizations were carried out using the “ggplot2” package (v3.5.1). Boxplots display the median and interquartile range. All analyses and genomic coordinates are based on the hg38 human genome assembly.

## Data availability

Sequences used are available at the NCBI SRA under accession number PRJNA1291813. All software new to this study is available at https://github.com/agmcfarland/BEiGUIDE and https://github.com/agmcfarland/CRISPRito.

## Acknowledgements

We are grateful to members of the Herbst-Nowrouzi and Bushman laboratories for help and suggestions. We thank Laurie Zimmerman for help with artwork. The authors thank Max Eldabbas, Emileigh Maddox, Tanishk Sinha, and Jiayi Shu of the Human Immunology Core at the Perelman School of Medicine at the University of Pennsylvania for assistance with providing isolated human normal T cell populations. The HIC is supported in part by NIH P30 AI045008 and P30 CA016520.

## Author Contributions

AGM, SKO, FDB, and FHN conceptualized the study. SKO, DT, EK, and NC carried out cell-based assays; AGM and FHN carried out bioinformatic analysis; EC and SR carried out DNA manipulations; all authors contributed ideas and materials; AGM, FDB, and FHN wrote the manuscript; all authors contributed edits and/or approved the manuscript.

## Supplementary Material

Supplementary Table 1. Published methods for monitoring off-target base editing.

Supplementary Table 2. The number of predicted off-target sites in iGUIDE experiments with relevant metadata. Both the total number of dsODN incorporations and in transcription unit (in_TU) are listed.

Supplementary Table 3. Metadata for all samples used in BEiGUIDE/iGUIDE and

rhAmpSeq experiments. The analysis column indicates what application the sample was used in. Accession lists the BioProject ID deposited under SRA ID PRJNA1291813.

Supplementary Table 4. Donors used in the study and marked for usage in iGUIDE/BEiGUIDE and/or rhAmpSeq experiments.

Supplementary Table 5. The top five shared potential off-targets identified in all seven normal T cell donors using iGUIDE. Aligned sequence and mismatch is in reference to PTPRC gRNA.

Supplementary Table 6. 14 potential off-target sites shared in at least three out of seven normal T cell donors. Aligned sequence and mismatch is in reference to PTPRC gRNA.

Supplementary Table 7. The number of predicted off-target sites for all cut-predicting bioinformatic tools. Both the total number of dsODN incorporations and in transcription unit (in_TU) are listed.

Supplementary Table 8. The genomic location of 144 targets selected for amplicon sequencing using rhAmpSeq.

Supplementary Table 9. An example of ranking off target edit sites using PTPRC data.

Supplementary 10. An example of choices of weights for ranking off target sites. Supplementary Table 11. All oligonucleotides used in study. If used previously, a citation is provided.

Supplementary Table 12. List of key reagents used in the study.

### Supplementary Figures

**Supplementary Figure 1.**
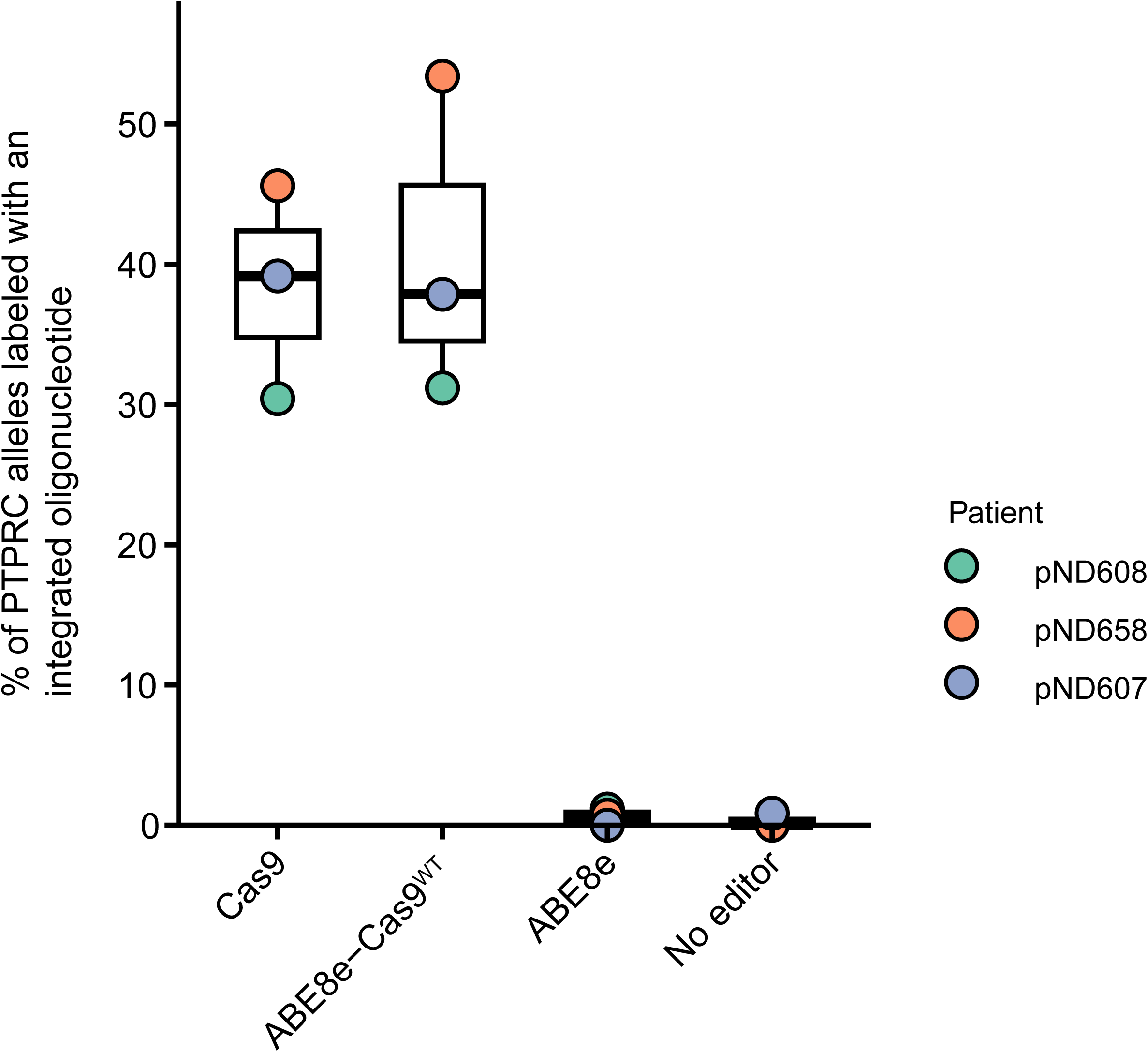
Comparison of marking efficiency over three cell donors (indicated by the three colors and key at right). The x-axis shows the enzyme used, the y-axis shows the efficiency of marking (% of oligonucleotide incorporation at *PTPRC* as a % of the total).

**Supplementary Figure 2.**
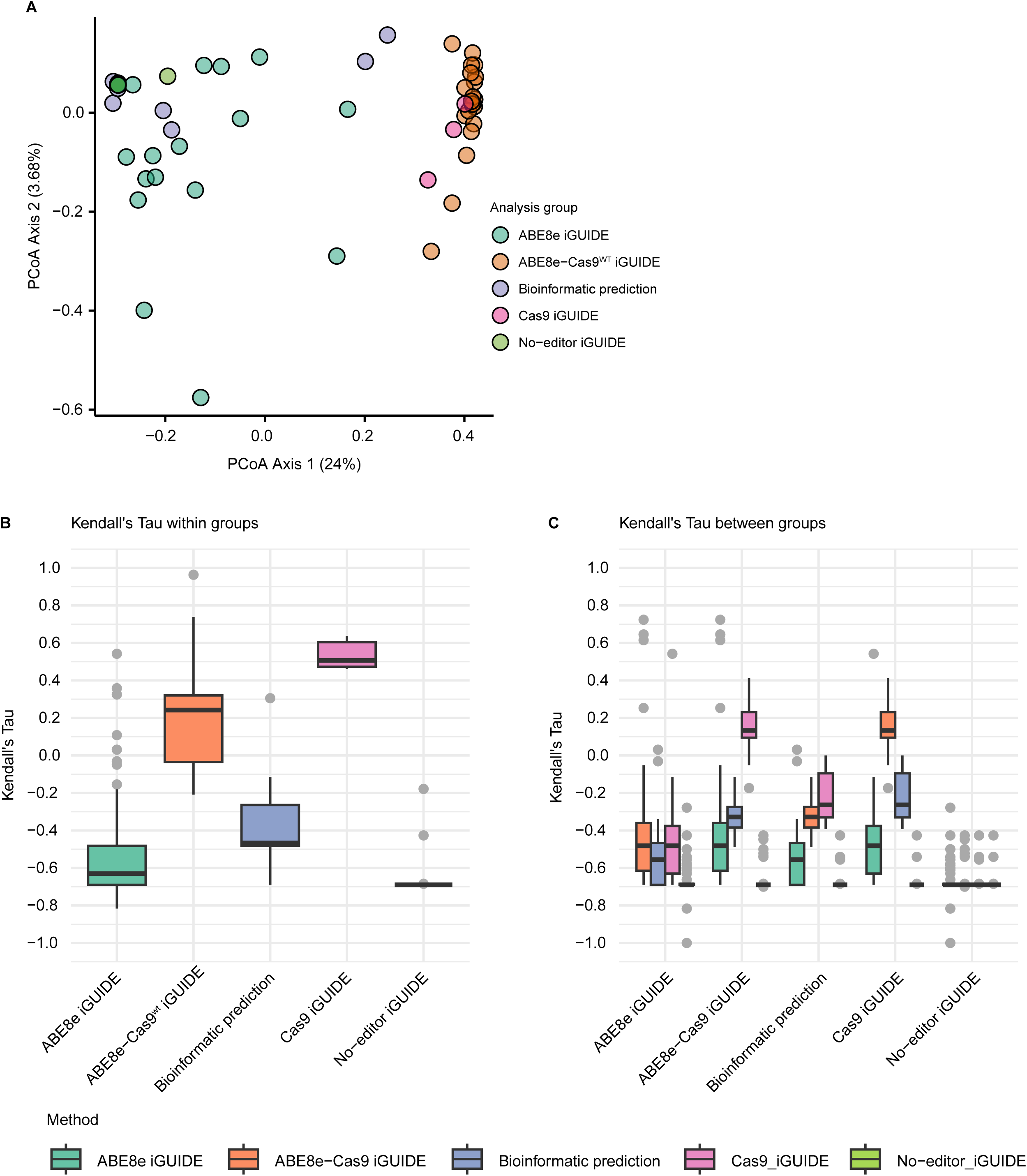
Comparison of off-target sites identified in the iGUIDE data and using bioinformatic methods. A) Principal coordinate analysis to compare the aggregate rankings of sites by each method. Both marking by oligonucleotide incorporation (iGUIDE) and bioinformatic methods are shown. The methods used are indicated by the color code to the right. The percent variance explained by each axis is shown in parenthesis. B) The within-group rank similarity of the top ten targets of *PTPRC* sgRNA, with method listed on the x-axis. Distribution is composed of biological replicates. Similarity was assessed using Kendall’s Tau. C) Between-group rank similarities, with pair-wise comparison of biological replicates in method group (x-axis) to biological replicates from method groups displayed in the legend.

**Supplementary Figure 3.**
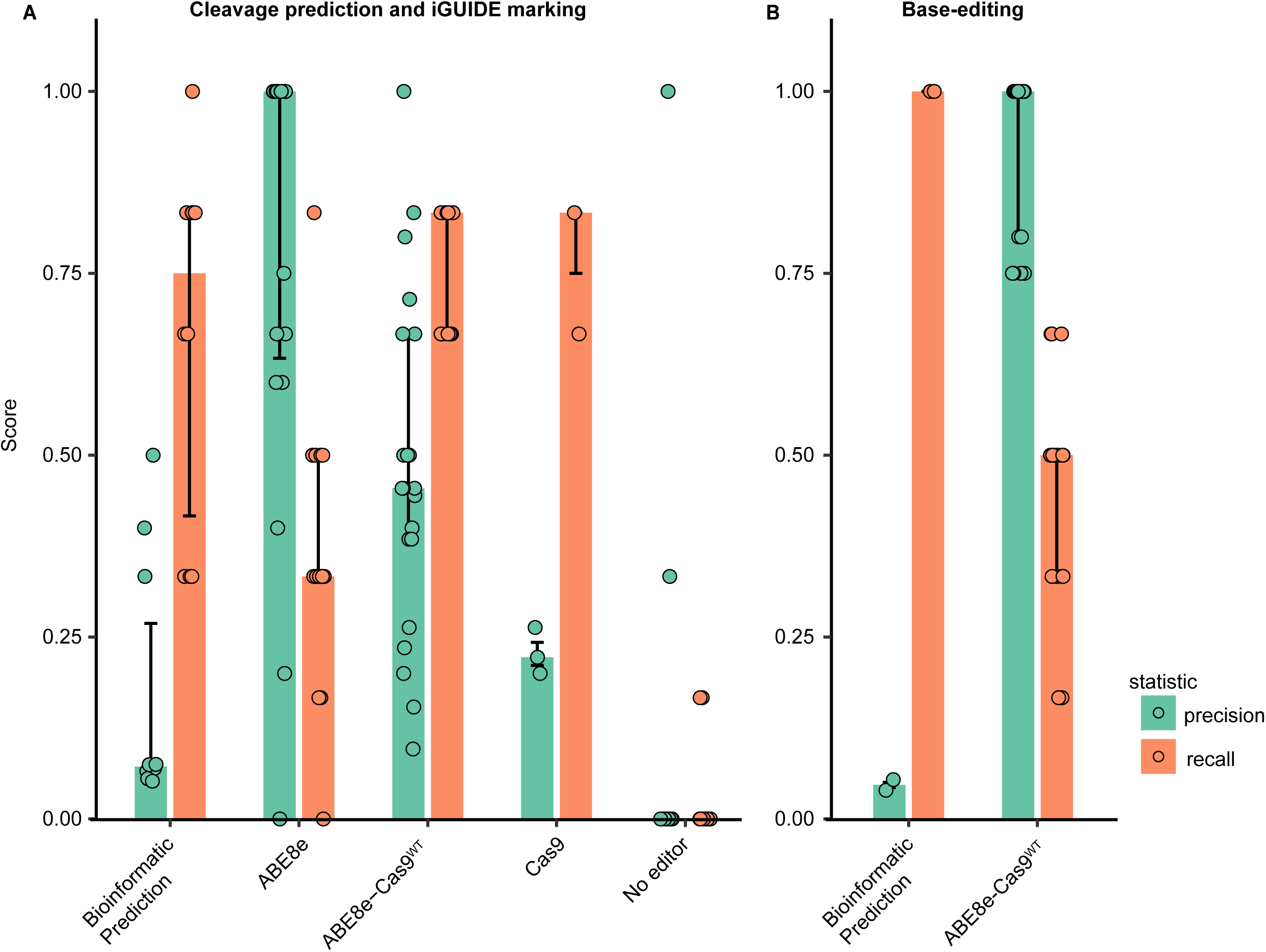
Comparison of methods for predicting sites of editing compared to results from actual ABE8e editing analyzed using the targeted amplicon panel. Results are compared by quantification of precision (true positives/true+false positives) and recall (true positives/true positives+false negatives) for the target sites analyzed by each method and compared to “truth” from the analysis of ABE8e and amplicon sequencing. A) Predictions for results of ABE8e editing using marking by oligonucleotide incorporation at dsDNA breaks (iGUIDE). “Computational” denotes the prediction of incorporation by the nine bioinformatic panels. “ABE8e” denotes comparison to the rare oligonucleotide incorporations detected in the presence of the ABE8e editor. “ABE8e-Cas9^WT^“ indicates comparison to sites of oligonucleotide incorporation in the presence of the ABE8e-Cas9^WT^ enzyme. “Cas9” denotes sites of oligonucleotide incorporation in the presence of the Cas9 editor. “No editor” indicates comparison to samples with no editor and so incorporation at background dsDNA breaks. B) Calls for sites of base editing compared to results from ABE8e and amplicon sequencing. “Computational” denotes pooled results of analysis from the two bioinformatic pipelines. “ABE8e-Cas9^WT^“ denotes analysis of editing at sites of oligonucleotide incorporation directed by the ABE8e-Cas9^WT^ enzyme.

**Supplementary Figure 4.**
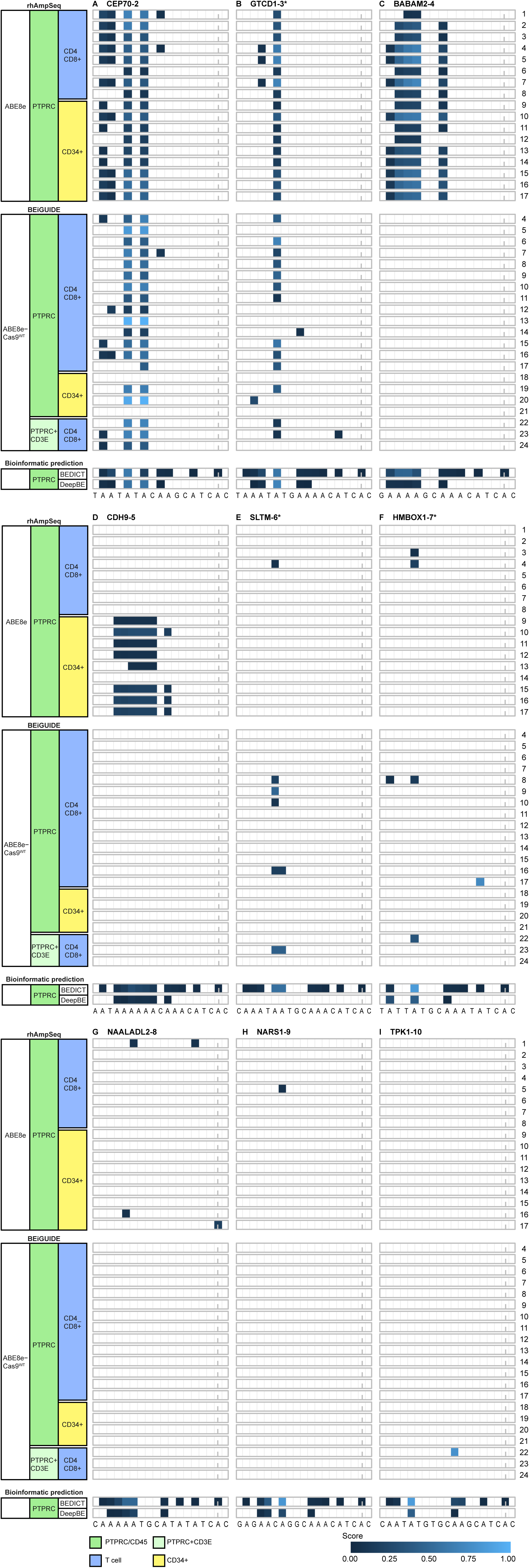
Comparison of bases edited at off target sites. A-I. Locations of edited bases at all off-target sites. The sequence of the matching. Samples are shown in the rows. A) Sites of base editing by the ABE8e-Cas9^WT^ at sites of oligonucleotide incorporation. B) Sites of base editing generated by the ABE8e base editing analyzed using the amplicon panel. C) Sites of base editing predicted by two bioinformatic tools. The extent of base editing is color-coded as indicated at the bottom. Numbering of the protospacer site starts on the left with “0”. The Cas9-targeted cut site is marked with the vertical dash. Asterisks and tildes indicate the site is in a transcription unit and in an oncogene, respectively.

