## Supplementary Table 2 for "An integrated enzymatic and computational pipeline for quantifying off-target base-editing"

**Supplementary Table 2.** **Summary of predicted off-target sites using iGUIDE-Seq.**

| **Sample** | **Patient** | **CellType** | **Timepoint** | **Enzyme** | **sgRNA** | **dsODN** | **total_hits** | **in_TU** |
| --- | --- | --- | --- | --- | --- | --- | --- | --- |
| GTSP6619 | pND608 | CD4+_CD8+ | d4 | Cas9 | PTPRC | 10 | 697 | 271 |
| GTSP6661 | pND658 | CD4+_CD8+ | d4 | Cas9 | PTPRC | 10 | 657 | 253 |
| GTSP6686 | pND607 | CD4+_CD8+ | d4 | Cas9 | PTPRC | 10 | 533 | 222 |
| GTSP6045 | p08718-032 | CD4+_CD8+ | d4 | ABE8e-Cas9 | PTPRC | 10 | 234 | 73 |
| GTSP5614 | pND567 | CD4+_CD8+ | d4 | ABE8e-Cas9 | PTPRC | 10 | 3389 | 1382 |
| GTSP5649 | p08718-039 | CD34+ | d4 | ABE8e-Cas9 | PTPRC | 10 | 940 | 365 |
| GTSP5619 | pTMP491 | CD4+_CD8+ | d4 | ABE8e-Cas9 | PTPRC | 10 | 222 | 92 |
| GTSP6618 | pND608 | CD4+_CD8+ | d4 | ABE8e-Cas9 | PTPRC | 10 | 733 | 246 |
| GTSP6685 | pND607 | CD4+_CD8+ | d4 | ABE8e-Cas9 | PTPRC | 10 | 492 | 173 |
| GTSP6249 | p08718-032 | CD34+ | d4 | ABE8e-Cas9 | PTPRC | 10 | 114 | 33 |
| GTSP6660 | pND658 | CD4+_CD8+ | d4 | ABE8e-Cas9 | PTPRC | 10 | 366 | 134 |
| GTSP6243 | p08718-043 | CD4+_CD8+ | d4 | ABE8e-Cas9 | PTPRC | 10 | 589 | 267 |
| GTSP5490 | pND579 | CD4+_CD8+ | d4 | ABE8e-Cas9 | PTPRC | 10 | 53 | 22 |
| GTSP6254 | p08718-043 | CD34+ | d4 | ABE8e-Cas9 | PTPRC | 10 | 166 | 54 |
| GTSP5613 | pND567 | CD4+_CD8+ | d4 | ABE8e-Cas9 | PTPRC | 1 | 197 | 90 |
| GTSP5648 | p08718-039 | CD34+ | d4 | ABE8e-Cas9 | PTPRC | 1 | 97 | 29 |
| GTSP5618 | pTMP491 | CD4+_CD8+ | d4 | ABE8e-Cas9 | PTPRC | 1 | 31 | 12 |
| GTSP5496 | pND579 | CD4+_CD8+ | d7 | ABE8e-Cas9 | PTPRC | 10 | 26 | 7 |
| GTSP5489 | pND579 | CD4+_CD8+ | d4 | ABE8e-Cas9 | PTPRC | 1 | 9 | 3 |
| GTSP5949 | p08718-039 | CD4+_CD8+ | d4 | ABE8e-Cas9 | PTPRC | 10 | 7 | 4 |
| GTSP5495 | pND579 | CD4+_CD8+ | d7 | ABE8e-Cas9 | PTPRC | 1 | 11 | 5 |
| GTSP6244 | p08718-043 | CD4+_CD8+ | d4 | ABE8e-Cas9 | PTPRC_CD3E | 10 | 739 | 283 |
| GTSP6046 | p08718-032 | CD4+_CD8+ | d4 | ABE8e-Cas9 | PTPRC_CD3E | 10 | 463 | 138 |
| GTSP5950 | p08718-039 | CD4+_CD8+ | d4 | ABE8e-Cas9 | PTPRC_CD3E | 10 | 70 | 27 |
| GTSP6684 | pND607 | CD4+_CD8+ | d4 | ABE8e | PTPRC | 10 | 1477 | 557 |
| GTSP5612 | pND567 | CD4+_CD8+ | d4 | ABE8e | PTPRC | 10 | 2917 | 1196 |
| GTSP6240 | p08718-043 | CD4+_CD8+ | d4 | ABE8e | PTPRC | 10 | 425 | 161 |
| GTSP5647 | p08718-039 | CD34+ | d4 | ABE8e | PTPRC | 10 | 587 | 205 |
| GTSP6042 | p08718-032 | CD4+_CD8+ | d4 | ABE8e | PTPRC | 10 | 131 | 35 |
| GTSP6617 | pND608 | CD4+_CD8+ | d4 | ABE8e | PTPRC | 10 | 381 | 129 |
| GTSP5946 | p08718-039 | CD4+_CD8+ | d4 | ABE8e | PTPRC | 10 | 112 | 30 |
| GTSP6659 | pND658 | CD4+_CD8+ | d4 | ABE8e | PTPRC | 10 | 222 | 84 |
| GTSP5488 | pND579 | CD4+_CD8+ | d4 | ABE8e | PTPRC | 10 | 91 | 31 |
| GTSP5617 | pTMP491 | CD4+_CD8+ | d4 | ABE8e | PTPRC | 10 | 130 | 57 |
| GTSP5487 | pND579 | CD4+_CD8+ | d4 | ABE8e | PTPRC | 1 | 5 | 3 |
| GTSP5494 | pND579 | CD4+_CD8+ | d7 | ABE8e | PTPRC | 10 | 18 | 9 |
| GTSP6248 | p08718-032 | CD34+ | d4 | ABE8e | PTPRC | 10 | 117 | 41 |
| GTSP6253 | p08718-043 | CD34+ | d4 | ABE8e | PTPRC | 10 | 64 | 16 |
| GTSP5493 | pND579 | CD4+_CD8+ | d7 | ABE8e | PTPRC | 1 | 2 | 1 |
| GTSP5646 | p08718-039 | CD34+ | d4 | ABE8e | PTPRC | 1 | 45 | 13 |
| GTSP6241 | p08718-043 | CD4+_CD8+ | d4 | ABE8e | PTPRC_CD3E | 10 | 891 | 364 |
| GTSP6043 | p08718-032 | CD4+_CD8+ | d4 | ABE8e | PTPRC_CD3E | 10 | 146 | 57 |
| GTSP5947 | p08718-039 | CD4+_CD8+ | d4 | ABE8e | PTPRC_CD3E | 10 | 1 | 0 |
| GTSP5611 | pND567 | CD4+_CD8+ | d4 | No-editor | PTPRC | 10 | 2598 | 1083 |
| GTSP6658 | pND658 | CD4+_CD8+ | d4 | No-editor | PTPRC | 10 | 291 | 102 |
| GTSP6616 | pND608 | CD4+_CD8+ | d4 | No-editor | PTPRC | 10 | 532 | 206 |
| GTSP5645 | p08718-039 | CD34+ | d4 | No-editor | PTPRC | 10 | 718 | 270 |
| GTSP5486 | pND579 | CD4+_CD8+ | d4 | No-editor | PTPRC | 10 | 70 | 28 |
| GTSP6683 | pND607 | CD4+_CD8+ | d4 | No-editor | PTPRC | 10 | 296 | 110 |
| GTSP6247 | p08718-032 | CD34+ | d4 | No-editor | PTPRC | 10 | 70 | 31 |
| GTSP6252 | p08718-043 | CD34+ | d4 | No-editor | PTPRC | 10 | 73 | 29 |
| GTSP5616 | pTMP491 | CD4+_CD8+ | d4 | No-editor | PTPRC | 10 | 152 | 45 |
| GTSP5644 | p08718-039 | CD34+ | d4 | No-editor | PTPRC | 1 | 37 | 18 |
| GTSP5492 | pND579 | CD4+_CD8+ | d7 | No-editor | PTPRC | 10 | 14 | 3 |
| GTSP5485 | pND579 | CD4+_CD8+ | d4 | No-editor | PTPRC | 1 | 7 | 2 |
| GTSP5491 | pND579 | CD4+_CD8+ | d7 | No-editor | PTPRC | 1 | 1 | 1 |
| GTSP5944 | p08718-039 | CD4+_CD8+ | d4 | No-editor | PTPRC_CD3E | 10 | 47 | 15 |
| GTSP6040 | p08718-032 | CD4+_CD8+ | d4 | No-editor | PTPRC_CD3E | 10 | 144 | 46 |
| GTSP6238 | p08718-043 | CD4+_CD8+ | d4 | No-editor | PTPRC_CD3E | 10 | 165 | 70 |
| GTSP6251 | p08718-043 | CD34+ | d4 | No-editor | No-sgRNA | 10 | 164 | 59 |
| GTSP6246 | p08718-032 | CD34+ | d4 | No-editor | No-sgRNA | 10 | 99 | 46 |
| GTSP5610 | pND567 | CD4+_CD8+ | d4 | No-editor | No-sgRNA | 10 | 2369 | 971 |
| GTSP5615 | pTMP491 | CD4+_CD8+ | d4 | No-editor | No-sgRNA | 10 | 108 | 37 |
