## Supplementary Table 3 for "An integrated enzymatic and computational pipeline for quantifying off-target base-editing"

**Supplementary Table 3. Metadata and accession numbers for sequence data sets used in this study (SRA ID PRJNA1291813)**

| **SpecimenAccNum** | **CellType** | **Patient** | **Timepoint** | **Enzyme** | **sgRNA** | **dsODN** | **analysis** | **Accession** |
| --- | --- | --- | --- | --- | --- | --- | --- | --- |
| GTSP5485 | CD4+_CD8+ | pND579 | d4 | No-editor | PTPRC | 1 | iGUIDE/BEiGUIDE | SAMN49978940 |
| GTSP5486 | CD4+_CD8+ | pND579 | d4 | No-editor | PTPRC | 10 | iGUIDE/BEiGUIDE | SAMN49978941 |
| GTSP5487 | CD4+_CD8+ | pND579 | d4 | ABE8e | PTPRC | 1 | iGUIDE/BEiGUIDE | SAMN49978942 |
| GTSP5488 | CD4+_CD8+ | pND579 | d4 | ABE8e | PTPRC | 10 | iGUIDE/BEiGUIDE | SAMN49978943 |
| GTSP5489 | CD4+_CD8+ | pND579 | d4 | ABE8e-Cas9 | PTPRC | 1 | iGUIDE/BEiGUIDE | SAMN49978944 |
| GTSP5490 | CD4+_CD8+ | pND579 | d4 | ABE8e-Cas9 | PTPRC | 10 | iGUIDE/BEiGUIDE | SAMN49978945 |
| GTSP5491 | CD4+_CD8+ | pND579 | d7 | No-editor | PTPRC | 1 | iGUIDE/BEiGUIDE | SAMN49978946 |
| GTSP5492 | CD4+_CD8+ | pND579 | d7 | No-editor | PTPRC | 10 | iGUIDE/BEiGUIDE | SAMN49978947 |
| GTSP5493 | CD4+_CD8+ | pND579 | d7 | ABE8e | PTPRC | 1 | iGUIDE/BEiGUIDE | SAMN49978948 |
| GTSP5494 | CD4+_CD8+ | pND579 | d7 | ABE8e | PTPRC | 10 | iGUIDE/BEiGUIDE | SAMN49978949 |
| GTSP5495 | CD4+_CD8+ | pND579 | d7 | ABE8e-Cas9 | PTPRC | 1 | iGUIDE/BEiGUIDE | SAMN49978950 |
| GTSP5496 | CD4+_CD8+ | pND579 | d7 | ABE8e-Cas9 | PTPRC | 10 | iGUIDE/BEiGUIDE | SAMN49978951 |
| GTSP5610 | CD4+_CD8+ | pND567 | d4 | No-editor | No-sgRNA | 10 | iGUIDE/BEiGUIDE | SAMN49978952 |
| GTSP5611 | CD4+_CD8+ | pND567 | d4 | No-editor | PTPRC | 10 | iGUIDE/BEiGUIDE | SAMN49978953 |
| GTSP5612 | CD4+_CD8+ | pND567 | d4 | ABE8e | PTPRC | 10 | iGUIDE/BEiGUIDE | SAMN49978954 |
| GTSP5613 | CD4+_CD8+ | pND567 | d4 | ABE8e-Cas9 | PTPRC | 1 | iGUIDE/BEiGUIDE | SAMN49978955 |
| GTSP5614 | CD4+_CD8+ | pND567 | d4 | ABE8e-Cas9 | PTPRC | 10 | iGUIDE/BEiGUIDE | SAMN49978956 |
| GTSP5615 | CD4+_CD8+ | pTMP491 | d4 | No-editor | No-sgRNA | 10 | iGUIDE/BEiGUIDE | SAMN49978957 |
| GTSP5616 | CD4+_CD8+ | pTMP491 | d4 | No-editor | PTPRC | 10 | iGUIDE/BEiGUIDE | SAMN49978958 |
| GTSP5617 | CD4+_CD8+ | pTMP491 | d4 | ABE8e | PTPRC | 10 | iGUIDE/BEiGUIDE | SAMN49978959 |
| GTSP5618 | CD4+_CD8+ | pTMP491 | d4 | ABE8e-Cas9 | PTPRC | 1 | iGUIDE/BEiGUIDE | SAMN49978960 |
| GTSP5619 | CD4+_CD8+ | pTMP491 | d4 | ABE8e-Cas9 | PTPRC | 10 | iGUIDE/BEiGUIDE | SAMN49978961 |
| GTSP5644 | CD34+ | p08718-039 | d4 | No-editor | PTPRC | 1 | iGUIDE/BEiGUIDE | SAMN49978962 |
| GTSP5645 | CD34+ | p08718-039 | d4 | No-editor | PTPRC | 10 | iGUIDE/BEiGUIDE | SAMN49978963 |
| GTSP5646 | CD34+ | p08718-039 | d4 | ABE8e | PTPRC | 1 | iGUIDE/BEiGUIDE | SAMN49978964 |
| GTSP5647 | CD34+ | p08718-039 | d4 | ABE8e | PTPRC | 10 | iGUIDE/BEiGUIDE | SAMN49978965 |
| GTSP5648 | CD34+ | p08718-039 | d4 | ABE8e-Cas9 | PTPRC | 1 | iGUIDE/BEiGUIDE | SAMN49978966 |
| GTSP5649 | CD34+ | p08718-039 | d4 | ABE8e-Cas9 | PTPRC | 10 | iGUIDE/BEiGUIDE | SAMN49978967 |
| GTSP5944 | CD4+_CD8+ | p08718-039 | d4 | No-editor | PTPRC_CD3E | 10 | iGUIDE/BEiGUIDE | SAMN49978968 |
| GTSP5946 | CD4+_CD8+ | p08718-039 | d4 | ABE8e | PTPRC | 10 | iGUIDE/BEiGUIDE | SAMN49978969 |
| GTSP5947 | CD4+_CD8+ | p08718-039 | d4 | ABE8e | PTPRC_CD3E | 10 | iGUIDE/BEiGUIDE | SAMN49978970 |
| GTSP5949 | CD4+_CD8+ | p08718-039 | d4 | ABE8e-Cas9 | PTPRC | 10 | iGUIDE/BEiGUIDE | SAMN49978971 |
| GTSP5950 | CD4+_CD8+ | p08718-039 | d4 | ABE8e-Cas9 | PTPRC_CD3E | 10 | iGUIDE/BEiGUIDE | SAMN49978972 |
| GTSP6040 | CD4+_CD8+ | p08718-032 | d4 | No-editor | PTPRC_CD3E | 10 | iGUIDE/BEiGUIDE | SAMN49978973 |
| GTSP6042 | CD4+_CD8+ | p08718-032 | d4 | ABE8e | PTPRC | 10 | iGUIDE/BEiGUIDE | SAMN49978974 |
| GTSP6043 | CD4+_CD8+ | p08718-032 | d4 | ABE8e | PTPRC_CD3E | 10 | iGUIDE/BEiGUIDE | SAMN49978975 |
| GTSP6045 | CD4+_CD8+ | p08718-032 | d4 | ABE8e-Cas9 | PTPRC | 10 | iGUIDE/BEiGUIDE | SAMN49978976 |
| GTSP6046 | CD4+_CD8+ | p08718-032 | d4 | ABE8e-Cas9 | PTPRC_CD3E | 10 | iGUIDE/BEiGUIDE | SAMN49978977 |
| GTSP6238 | CD4+_CD8+ | p08718-043 | d4 | No-editor | PTPRC_CD3E | 10 | iGUIDE/BEiGUIDE | SAMN49978978 |
| GTSP6240 | CD4+_CD8+ | p08718-043 | d4 | ABE8e | PTPRC | 10 | iGUIDE/BEiGUIDE | SAMN49978979 |
| GTSP6241 | CD4+_CD8+ | p08718-043 | d4 | ABE8e | PTPRC_CD3E | 10 | iGUIDE/BEiGUIDE | SAMN49978980 |
| GTSP6243 | CD4+_CD8+ | p08718-043 | d4 | ABE8e-Cas9 | PTPRC | 10 | iGUIDE/BEiGUIDE | SAMN49978981 |
| GTSP6244 | CD4+_CD8+ | p08718-043 | d4 | ABE8e-Cas9 | PTPRC_CD3E | 10 | iGUIDE/BEiGUIDE | SAMN49978982 |
| GTSP6247 | CD34+ | p08718-032 | d4 | No-editor | PTPRC | 10 | iGUIDE/BEiGUIDE | SAMN49978983 |
| GTSP6248 | CD34+ | p08718-032 | d4 | ABE8e | PTPRC | 10 | iGUIDE/BEiGUIDE | SAMN49978984 |
| GTSP6249 | CD34+ | p08718-032 | d4 | ABE8e-Cas9 | PTPRC | 10 | iGUIDE/BEiGUIDE | SAMN49978985 |
| GTSP6250 | CD34+ | p08718-043 | d4 | No-editor | No-sgRNA | 0 | iGUIDE/BEiGUIDE | SAMN49978986 |
| GTSP6251 | CD34+ | p08718-043 | d4 | No-editor | No-sgRNA | 10 | iGUIDE/BEiGUIDE | SAMN49978987 |
| GTSP6252 | CD34+ | p08718-043 | d4 | No-editor | PTPRC | 10 | iGUIDE/BEiGUIDE | SAMN49978988 |
| GTSP6253 | CD34+ | p08718-043 | d4 | ABE8e | PTPRC | 10 | iGUIDE/BEiGUIDE | SAMN49978989 |
| GTSP6254 | CD34+ | p08718-043 | d4 | ABE8e-Cas9 | PTPRC | 10 | iGUIDE/BEiGUIDE | SAMN49978990 |
| GTSP6616 | CD4+_CD8+ | pND608 | d4 | No-editor | PTPRC | 10 | iGUIDE/BEiGUIDE | SAMN49978991 |
| GTSP6617 | CD4+_CD8+ | pND608 | d4 | ABE8e | PTPRC | 10 | iGUIDE/BEiGUIDE | SAMN49978992 |
| GTSP6618 | CD4+_CD8+ | pND608 | d4 | ABE8e-Cas9 | PTPRC | 10 | iGUIDE/BEiGUIDE | SAMN49978993 |
| GTSP6619 | CD4+_CD8+ | pND608 | d4 | Cas9 | PTPRC | 10 | iGUIDE/BEiGUIDE | SAMN49978994 |
| GTSP6658 | CD4+_CD8+ | pND658 | d4 | No-editor | PTPRC | 10 | iGUIDE/BEiGUIDE | SAMN49978995 |
| GTSP6659 | CD4+_CD8+ | pND658 | d4 | ABE8e | PTPRC | 10 | iGUIDE/BEiGUIDE | SAMN49978996 |
| GTSP6660 | CD4+_CD8+ | pND658 | d4 | ABE8e-Cas9 | PTPRC | 10 | iGUIDE/BEiGUIDE | SAMN49978997 |
| GTSP6661 | CD4+_CD8+ | pND658 | d4 | Cas9 | PTPRC | 10 | iGUIDE/BEiGUIDE | SAMN49978998 |
| GTSP6683 | CD4+_CD8+ | pND607 | d4 | No-editor | PTPRC | 10 | iGUIDE/BEiGUIDE | SAMN49978999 |
| GTSP6684 | CD4+_CD8+ | pND607 | d4 | ABE8e | PTPRC | 10 | iGUIDE/BEiGUIDE | SAMN49979000 |
| GTSP6685 | CD4+_CD8+ | pND607 | d4 | ABE8e-Cas9 | PTPRC | 10 | iGUIDE/BEiGUIDE | SAMN49979001 |
| GTSP6686 | CD4+_CD8+ | pND607 | d4 | Cas9 | PTPRC | 10 | iGUIDE/BEiGUIDE | SAMN49979002 |
| HTS_SO81_01_S1 | CD4+_CD8+ | pND608 | d0 | No-editor | No-sgRNA | NA | rhAmp | SAMN49979003 |
| HTS_SO81_02_S2 | CD4+_CD8+ | pND608 | d0 | No-editor | PTPRC | NA | rhAmp | SAMN49979004 |
| HTS_SO81_03_S3 | CD4+_CD8+ | pND608 | d0 | ABE8e | PTPRC | NA | rhAmp | SAMN49979005 |
| HTS_SO81_05_S5 | CD4+_CD8+ | pND578 | d0 | No-editor | No-sgRNA | NA | rhAmp | SAMN49979006 |
| HTS_SO81_06_S6 | CD4+_CD8+ | pND578 | d0 | No-editor | PTPRC | NA | rhAmp | SAMN49979007 |
| HTS_SO81_07_S7 | CD4+_CD8+ | pND578 | d0 | ABE8e | PTPRC | NA | rhAmp | SAMN49979008 |
| HTS_SO81_10_S10 | CD4+_CD8+ | pND500 | d0 | No-editor | No-sgRNA | NA | rhAmp | SAMN49979009 |
| HTS_SO81_11_S11 | CD4+_CD8+ | pND500 | d0 | No-editor | No-sgRNA | NA | rhAmp | SAMN49979010 |
| HTS_SO81_12_S12 | CD4+_CD8+ | pND500 | d0 | No-editor | PTPRC | NA | rhAmp | SAMN49979011 |
| HTS_SO81_13_S13 | CD4+_CD8+ | pND500 | d0 | ABE8e | PTPRC | NA | rhAmp | SAMN49979012 |
| HTS_SO81_14_S14 | CD4+_CD8+ | pND500 | d0 | ABE8e | PTPRC | NA | rhAmp | SAMN49979013 |
| HTS_SO81_52_S52 | CD4+_CD8+ | pND627 | d0 | ABE8e | PTPRC | NA | rhAmp | SAMN49979014 |
| HTS_SO81_53_S53 | CD4+_CD8+ | pND627 | d0 | ABE8e | PTPRC | NA | rhAmp | SAMN49979015 |
| HTS_SO81_54_S54 | CD4+_CD8+ | pND627 | d0 | ABE8e | PTPRC | NA | rhAmp | SAMN49979016 |
| HTS_SO81_55_S55 | CD4+_CD8+ | pND627 | d0 | ABE8e | PTPRC | NA | rhAmp | SAMN49979017 |
| HTS_SO81_15_S15 | CD34+ | pND627 | d0 | No-editor | No-sgRNA | NA | rhAmp | SAMN49979018 |
| HTS_SO81_16_S16 | CD34+ | pND627 | d0 | No-editor | PTPRC | NA | rhAmp | SAMN49979019 |
| HTS_SO81_17_S17 | CD34+ | pND627 | d0 | ABE8e | PTPRC | NA | rhAmp | SAMN49979020 |
| HTS_SO81_21_S21 | CD34+ | p08718-032 | d0 | No-editor | No-sgRNA | NA | rhAmp | SAMN49979021 |
| HTS_SO81_22_S22 | CD34+ | p08718-032 | d0 | No-editor | PTPRC | NA | rhAmp | SAMN49979022 |
| HTS_SO81_23_S23 | CD34+ | p08718-032 | d0 | ABE8e | PTPRC | NA | rhAmp | SAMN49979023 |
| HTS_SO81_36_S36 | CD34+ | p08718-032 | d0 | No-editor | No-sgRNA | NA | rhAmp | SAMN49979024 |
| HTS_SO81_37_S37 | CD34+ | p08718-032 | d0 | No-editor | PTPRC | NA | rhAmp | SAMN49979025 |
| HTS_SO81_38_S38 | CD34+ | p08718-032 | d0 | ABE8e | PTPRC | NA | rhAmp | SAMN49979026 |
| HTS_SO81_45_S45 | CD34+ | p08718-032 | d0 | ABE8e | PTPRC | NA | rhAmp | SAMN49979027 |
| HTS_SO81_46_S46 | CD34+ | p08718-032 | d0 | ABE8e | PTPRC | NA | rhAmp | SAMN49979028 |
| HTS_SO81_47_S47 | CD34+ | p08718-032 | d0 | ABE8e | PTPRC | NA | rhAmp | SAMN49979029 |
| HTS_SO81_48_S48 | CD34+ | p08718-032 | d0 | ABE8e | PTPRC | NA | rhAmp | SAMN49979030 |
| HTS_SO81_49_S49 | CD34+ | p08718-032 | d0 | ABE8e | PTPRC | NA | rhAmp | SAMN49979031 |
| HTS_SO81_50_S50 | CD34+ | p08718-032 | d0 | ABE8e | PTPRC | NA | rhAmp | SAMN49979032 |
| HTS_SO81_51_S51 | CD34+ | p08718-032 | d0 | No-editor | PTPRC | NA | rhAmp | SAMN49979033 |
