## Supplementary Table 4 for "An integrated enzymatic and computational pipeline for quantifying off-target base-editing"

**Supplementary Table 4. Summary of donor samples used in the different experiments**

| **Donor** | **iGUIDE/BEiGUIDE** | **rhAmp-Seq** |
| --- | --- | --- |
| pND579 | 1 | NA |
| pND567 | 1 | NA |
| pTMP491 | 1 | NA |
| p08718-039 | 1 | NA |
| p08718-032 | 1 | 1 |
| p08718-043 | 1 | NA |
| pND608 | 1 | 1 |
| pND658 | 1 | NA |
| pND607 | 1 | NA |
| pND578 | NA | 1 |
| pND500 | NA | 1 |
| pND627 | NA | 1 |
