## Supplementary Table 5 for "An integrated enzymatic and computational pipeline for quantifying off-target base-editing"

**Supplementary Table 5.** **Top five shared potential off-target sites identified in seven normal T cell donors** (iGUIDE | cellular approach; * = within transcription unit, ~ = cancer associated gene)

| **Target** | **Gene ID** | **Edit Site** | **Aligned Sequence** | **Mismatch** | **Rel. abundance** | **Std Dev.** |
| --- | --- | --- | --- | --- | --- | --- |
| On-target | PTPRC*~ | chr1:+:198706754 | ....................T.. | 0 | 1 |  |
| Off#1 | CEP70 | chr3:+:138494328 | GT.....A...G........A.. | 4 | 0.367143 | 0.270044 |
| Off#2 | GTDC1* | chr2:-:143961596 | T.......A...........A.. | 2 | 0.131429 | 0.132844 |
| Off#3 | TPK1 | chr7:+:144862724 | ..T..G.....G........A.. | 3 | 0.029571 | 0.032928 |
| Off#4 | HMBOX1* | chr8:+:28930554 | GT.T........T.......T.. | 4 | 0.027571 | 0.029966 |
| Off#5 | KALRN* | chr3:-:124196101 | .C.T.G..............A.. | 3 | 0.021429 | 0.016762 |
