## Supplementary Table 6 for "An integrated enzymatic and computational pipeline for quantifying off-target base-editing"

**Supplementary Table 6.** **14 potential off-target sites shared in at least three out of seven normal T cell donors** (iGUIDE | cellular approach; * = within transcription unit, ~ = cancer associated gene).

| **Target** | **Gene ID** | **Edit Site** | **Aligned Sequence** | **Mismatch** | **Donors** | **Rel. abund.** |
| --- | --- | --- | --- | --- | --- | --- |
| Off#6 | NALF1* | chr13:+:107721971 | .T..ATAAG...........A.. | 6 | 6/7 | <0.01 |
| Off#7 | NR3C2 | chr4:-:148700496 | TG...G..T...........G.. | 4 | 5/7 |  |
| Off#8 | RAB6B | chr3:-:133903435 | .....TATA...........A.. | 4 |  |  |
| Off#9 | SYT8 | chr11:-:1825884 | .CTG.G....G.........C.. | 5 |  |  |
| Off#10 | THADA*~ | chr2:-:43508377 | T..G..G...G.........T.. | 4 |  |  |
| Off#11 | LOC100505549* | chr18:+:57630540 | .G..C.G.............C.. | 3 |  |  |
| Off#12 | CWH43 | chr4:-:50273364 | GTGGA.............A.A.A | 6 |  |  |
| Off#13 | BABAM2 | chr2:-:28359447 | CTG.A.A.............G.. | 5 | 4/7 |  |
| Off#14 | RBSN* | chr3:+:15096474 | .....G..A...........GA. | 2 |  |  |
| Off#15 | ZNF805* | chr19:-:57248979 | CTC....A.....T......G.. | 5 |  |  |
| Off#16 | VAPA | chr18:+:10074654 | CTGCAG..............G.. | 6 |  |  |
| Off#17 | MACROD2* | chr20:-:14311815 | ..G.......T...GGC.G.GT. | 6 |  |  |
| Off#18 | FRG1CP | chr20:-:28576421 | ...T....T...A..A.T.AAT. | 6 | 3/7 |  |
| Off#19 | DPYD* | chr1:+:97377036 | ....A.G.A........AGTCA. | 6 |  |  |
