## Supplementary Table 7 for "An integrated enzymatic and computational pipeline for quantifying off-target base-editing"

**Supplementary Table 7. Summary of predicted off-target sites for BE8 gRNA Cas9 cleavage using 10 different in silico tools**

| **method** | **total_hits** | **in_TU** |
| --- | --- | --- |
| ABEdeepoff | 745 | 308 |
| CasOFFinder | 745 | 308 |
| CCTop | 322 | 117 |
| CHOPCHOP | 44 | 18 |
| COSMID | 11 | 6 |
| CRISPOR | 808 | 259 |
| CRISPRme | 968 | 384 |
| CRISPRoff | 50 | 6 |
| CRISTA | 3181 | 1234 |
| IDT | 96 | 36 |
