## Supplementary Table 8 for "An integrated enzymatic and computational pipeline for quantifying off-target base-editing"

**Supplementary Table 8.** **Genomic locations selected for targeted amplicon sequencing**

| **Chromosome** | **Start** | **End** | **Assay ID** |
| --- | --- | --- | --- |
| chr3 | 31336119 | 31336291 | 1035RH.619A5FCAD142492Z0Z |
| chr22 | 33836983 | 33837223 | 1127RH.65FCEE86560D483Z0Z |
| chr18 | 33998150 | 33998335 | 1132RH.7DD0DDABBB284AEZ0Z |
| chr3 | 34875465 | 34875678 | 1165RH.B3A11FD5CD5E484Z0Z |
| chr5 | 37129899 | 37130068 | 1235RH.9EC04E8C04B143AZ0Z |
| chrX | 39004812 | 39005053 | 1297RH.7699FF425252411Z0Z |
| chr9 | 40018494 | 40018741 | 1328RH.160A09D64B5E485Z0Z |
| chr14 | 40300809 | 40300980 | 1333RH.15B9A62EB1DE41BZ0Z |
| chr2 | 41103488 | 41103708 | 1370RH.0D4AA7A212C44EFZ0Z |
| chr2 | 43508331 | 43508490 | 1449RH.D06DA35D5C6F4E9Z0Z |
| chr4 | 6235308 | 6235525 | 146RH.EF08CEB6926C4B4Z0Z |
| chr10 | 45109620 | 45109791 | 1510RH.05CAC48FC8C64C3Z0Z |
| chr13 | 45449330 | 45449513 | 1523RH.BE9A32AAC157493Z0Z |
| chr5 | 46181245 | 46181416 | 1546RH.D5EC6A5AE8E34A3Z0Z |
| chr19 | 46281259 | 46281429 | 1548RH.1C1C8FA5B72B4AEZ0Z |
| chr16 | 47235488 | 47235652 | 1579RH.E481C4E35AB345CZ0Z |
| chr22 | 49079706 | 49079935 | 1623RH.B099A99EF7824DBZ0Z |
| chr15 | 49245285 | 49245499 | 1629RH.F11445AE801E48FZ0Z |
| chr19 | 6749343 | 6749526 | 171RH.1CC31CF2C38142FZ0Z |
| chr10 | 50467732 | 50467919 | 1743RH.1F6AF979BBC949CZ0Z |
| chr5 | 51027172 | 51027364 | 1810RH.29EB32805946469Z0Z |
| chr3 | 51035952 | 51036133 | 1812RH.5A3E6971BEE5410Z0Z |
| chr7 | 1148427 | 1148659 | 18RH.628EE7DAF33746BZ0Z |
| chr4 | 52935402 | 52935596 | 1943RH.8D7AE6B5D4E443BZ0Z |
| chr14 | 53495795 | 53495969 | 1961RH.BB46480D41A4477Z0Z |
| chr7 | 7567825 | 7568033 | 196RH.FCA4EB067E0A4A8Z0Z |
| chr18 | 7811039 | 7811216 | 203RH.6A6189F138804E9Z0Z |
| chr6 | 7937051 | 7937217 | 207RH.7C0359D6CFCC4F8Z0Z |
| chr8 | 57472276 | 57472459 | 2095RH.A22CBEC7D912400Z0Z |
| chr18 | 57630447 | 57630650 | 2098RH.03DE457D2EED44DZ0Z |
| chr18 | 57695775 | 57695942 | 2099RH.CDF8A44FCF0C4F6Z0Z |
| chr15 | 58902074 | 58902282 | 2129RH.50311ABEA42E44BZ0Z |
| chr3 | 62540986 | 62541230 | 2246RH.E7C25A8C19B0486Z0Z |
| chr11 | 63312819 | 63312986 | 2276RH.B7161689C38445CZ0Z |
| chr8 | 8749091 | 8749272 | 227RH.DB8EF3336C214BEZ0Z |
| chr4 | 63398124 | 63398344 | 2281RH.0D05689F6B49481Z0Z |
| chrX | 65186820 | 65187016 | 2342RH.36ACF955B8E0432Z0Z |
| chr17 | 66358639 | 66358823 | 2397RH.53A8F6FB1C794A1Z0Z |
| chr1 | 66705399 | 66705580 | 2403RH.0095D21FE536427Z0Z |
| chrX | 68428696 | 68428938 | 2448RH.E0431F27FF704A1Z0Z |
| chr11 | 69127664 | 69127873 | 2468RH.82C2D3022E5B44AZ0Z |
| chr15 | 71165994 | 71166211 | 2548RH.287107F01D5C4BEZ0Z |
| chr13 | 72141477 | 72141649 | 2578RH.84617C8D0A83486Z0Z |
| chr18 | 72164867 | 72165061 | 2579RH.F58D39C4BBCE4ABZ0Z |
| chr14 | 72356398 | 72356578 | 2587RH.2EA5CA8A765B4F4Z0Z |
| chr4 | 73940556 | 73940736 | 2637RH.DC1BCA3CF998407Z0Z |
| chr4 | 74033674 | 74033857 | 2640RH.8ADD3DCDF6CE4FBZ0Z |
| chr13 | 74081245 | 74081478 | 2643RH.CE545E8290F04AFZ0Z |
| chr9 | 74628441 | 74628609 | 2655RH.B3FD1D054605492Z0Z |
| chr8 | 74745504 | 74745735 | 2657RH.558F5D87F5FC4B6Z0Z |
| chr14 | 76697668 | 76697906 | 2721RH.B7BF3596A0F3484Z0Z |
| chr10 | 76797026 | 76797219 | 2726RH.D26D71B343C649EZ0Z |
| chr11 | 78443373 | 78443543 | 2790RH.CA2F4E7BDBFB4E7Z0Z |
| chr14 | 78506224 | 78506453 | 2792RH.0AB5694CB306478Z0Z |
| chr18 | 10074576 | 10074747 | 280RH.F47FFA919A69426Z0Z |
| chr4 | 79575854 | 79576097 | 2828RH.CE86C1125ADF435Z0Z |
| chr17 | 79787685 | 79787853 | 2836RH.87CA33564CA742AZ0Z |
| chr7 | 81063146 | 81063339 | 2880RH.99130009F06942FZ0Z |
| chr5 | 81289661 | 81289850 | 2889RH.8CF58B40ACAE431Z0Z |
| chr4 | 83215654 | 83215825 | 2962RH.5B3C12D3F2084B5Z0Z |
| chr8 | 84258719 | 84258882 | 2995RH.E2D7AD2B87ED49FZ0Z |
| chr10 | 85082268 | 85082504 | 3013RH.ABE6F7DD63DF43EZ0Z |
| chr1 | 85158529 | 85158772 | 3015RH.C2BD8FC764B340FZ0Z |
| chr9 | 86538711 | 86538899 | 3057RH.8B95F98E717141CZ0Z |
| chr1 | 88227656 | 88227876 | 3106RH.5861DA747E834C3Z0Z |
| chr2 | 89618186 | 89618430 | 3149RH.95797A949BF64C1Z0Z |
| chr14 | 89631236 | 89631416 | 3151RH.B5D86BEF3F2D40CZ0Z |
| chr6 | 92046170 | 92046353 | 3206RH.48D93D137D5742CZ0Z |
| chr6 | 92243745 | 92243976 | 3210RH.C6340DB44C064B1Z0Z |
| chr11 | 92297807 | 92297978 | 3212RH.A96B72D181134D7Z0Z |
| chr15 | 94599983 | 94600154 | 3268RH.1C552DBBFA564E6Z0Z |
| chr2 | 1.03E+08 | 1.03E+08 | 3492RH.480142B7FE9E407Z0Z |
| chrX | 1.04E+08 | 1.04E+08 | 3499RH.73A985EFEC98494Z0Z |
| chr12 | 1.04E+08 | 1.04E+08 | 3514RH.DAE07D27A3FB475Z0Z |
| chr1 | 1.04E+08 | 1.04E+08 | 3516RH.37C744AAF832486Z0Z |
| chr6 | 1.08E+08 | 1.08E+08 | 3621RH.D649A0EC1EA144EZ0Z |
| chr10 | 1.08E+08 | 1.08E+08 | 3627RH.D4D42A2881AF477Z0Z |
| chr9 | 1.11E+08 | 1.11E+08 | 3705RH.C1AB424E8DB743DZ0Z |
| chr4 | 1.12E+08 | 1.12E+08 | 3714RH.774202BF9809441Z0Z |
| chr12 | 1.13E+08 | 1.13E+08 | 3743RH.769AE69EA0314FAZ0Z |
| chrX | 1.13E+08 | 1.13E+08 | 3754RH.34E9E3D6ABE3439Z0Z |
| chr12 | 1.13E+08 | 1.13E+08 | 3764RH.6DF434CE46AC43AZ0Z |
| chr8 | 1.15E+08 | 1.15E+08 | 3803RH.DBF8DDEE9DA8409Z0Z |
| chr7 | 1.15E+08 | 1.15E+08 | 3809RH.636FF6012CEE455Z0Z |
| chr4 | 1.16E+08 | 1.16E+08 | 3820RH.96B434E156EB4EFZ0Z |
| chr12 | 1.19E+08 | 1.19E+08 | 3880RH.5EC0DDABB50941BZ0Z |
| chr7 | 1.19E+08 | 1.19E+08 | 3881RH.67CCB51C28C34E5Z0Z |
| chr5 | 1.2E+08 | 1.2E+08 | 3900RH.8354E1EEBC48491Z0Z |
| chr1 | 1.21E+08 | 1.21E+08 | 3921RH.A40A2496959D460Z0Z |
| chr11 | 1.22E+08 | 1.22E+08 | 3940RH.ADB8B744D6DB48BZ0Z |
| chr5 | 1.24E+08 | 1.24E+08 | 3976RH.DF4E53E4DD5A488Z0Z |
| chr3 | 1.24E+08 | 1.24E+08 | 3978RH.8E8DA1A292BA48AZ0Z |
| chr3 | 1.25E+08 | 1.25E+08 | 3994RH.6B7DE361066F437Z0Z |
| chr3 | 1.26E+08 | 1.26E+08 | 4005RH.B84A4227E5B442BZ0Z |
| chr12 | 1.26E+08 | 1.26E+08 | 4012RH.A827BDD68C67422Z0Z |
| chr3 | 1.27E+08 | 1.27E+08 | 4030RH.E282E6FE0C39435Z0Z |
| chr5 | 1.31E+08 | 1.31E+08 | 4097RH.AB810418B950435Z0Z |
| chr8 | 1.32E+08 | 1.32E+08 | 4117RH.9C244683DFC3457Z0Z |
| chr3 | 1.35E+08 | 1.35E+08 | 4175RH.0693CA923BB84C5Z0Z |
| chr8 | 1.37E+08 | 1.37E+08 | 4194RH.C7160920023B476Z0Z |
| chr3 | 1.38E+08 | 1.38E+08 | 4217RH.15F5B9E789134B5Z0Z |
| chrX | 1.41E+08 | 1.41E+08 | 4246RH.8972152771F542EZ0Z |
| chr2 | 1.42E+08 | 1.42E+08 | 4255RH.8C62FDCC50CA41CZ0Z |
| chr2 | 1.44E+08 | 1.44E+08 | 4276RH.60E57232333B486Z0Z |
| chr7 | 1.45E+08 | 1.45E+08 | 4285RH.9B46BDF5672546CZ0Z |
| chr4 | 1.45E+08 | 1.45E+08 | 4299RH.B2EBAFBBD80B42FZ0Z |
| chr4 | 1.49E+08 | 1.49E+08 | 4343RH.2D2CA876FDF040CZ0Z |
| chr2 | 1.52E+08 | 1.52E+08 | 4377RH.7EB1158DD4C040CZ0Z |
| chr2 | 1.54E+08 | 1.54E+08 | 4403RH.9D46868552C4478Z0Z |
| chrX | 1.55E+08 | 1.55E+08 | 4409RH.9B0D233AF2EB4F1Z0Z |
| chr2 | 1.56E+08 | 1.56E+08 | 4418RH.F27B9D86252748EZ0Z |
| chr6 | 1.65E+08 | 1.65E+08 | 4527RH.575C39523B52496Z0Z |
| chr4 | 1.75E+08 | 1.75E+08 | 4620RH.34C8184F57E1474Z0Z |
| chr3 | 1.76E+08 | 1.76E+08 | 4626RH.C9B4EF959F254BAZ0Z |
| chr1 | 1.76E+08 | 1.76E+08 | 4630RH.7A6AB80815BC47FZ0Z |
| chr3 | 1.77E+08 | 1.77E+08 | 4635RH.9D21867198C94C1Z0Z |
| chr1 | 1.84E+08 | 1.84E+08 | 4692RH.FE213F00AA334ECZ0Z |
| chr1 | 1.99E+08 | 1.99E+08 | 4767RH.225982A3485E4AFZ0Z |
| chr1 | 2.02E+08 | 2.02E+08 | 4775RH.9B4D970E37734F2Z0Z |
| chr2 | 2.1E+08 | 2.1E+08 | 4801RH.6709343222BF4DCZ0Z |
| chr1 | 2.21E+08 | 2.21E+08 | 4837RH.1B157F773138479Z0Z |
| chr1 | 2.27E+08 | 2.27E+08 | 4857RH.39553ACF9027414Z0Z |
| chr1 | 2.41E+08 | 2.41E+08 | 4899RH.3BC2B0DEDE31443Z0Z |
| chr1 | 2.48E+08 | 2.48E+08 | 4914RH.ED2EF2FA003941FZ0Z |
| chr5 | 17245246 | 17245419 | 519RH.E2BC96B4FDF34B6Z0Z |
| chr16 | 17697633 | 17697825 | 529RH.DB4550BB2B26455Z0Z |
| chr10 | 17888324 | 17888494 | 537RH.F75793E8BCB3480Z0Z |
| chr3 | 19198643 | 19198824 | 600RH.AF7F645E6E08458Z0Z |
| chr19 | 20699886 | 20700051 | 644RH.18B65022731749BZ0Z |
| chr1 | 21826915 | 21827133 | 680RH.6205A0453454441Z0Z |
| chr7 | 22170919 | 22171135 | 692RH.67B3E12FFC71485Z0Z |
| chr1 | 22293185 | 22293356 | 700RH.AD74507F9F4C44EZ0Z |
| chr12 | 23018354 | 23018570 | 727RH.E5AA73B41EB6418Z0Z |
| chr3 | 23158212 | 23158401 | 733RH.12F332D92163403Z0Z |
| chr4 | 23344507 | 23344669 | 737RH.265A959074714CBZ0Z |
| chr21 | 25444487 | 25444660 | 828RH.718296D8B3274B7Z0Z |
| chr14 | 25602457 | 25602696 | 833RH.64D6C9450925453Z0Z |
| chr6 | 25709198 | 25709377 | 839RH.624CFF19C129415Z0Z |
| chr5 | 27172204 | 27172373 | 886RH.FA63720C0BAD4A4Z0Z |
| chr10 | 4123381 | 4123554 | 89RH.B6C6C38E9BEB4C2Z0Z |
| chr2 | 28359330 | 28359520 | 935RH.78DE8C3E5CE1421Z0Z |
| chr14 | 28807555 | 28807723 | 948RH.C5FFE160D89C4F5Z0Z |
| chr8 | 28930453 | 28930634 | 956RH.E9DFD67921DC4DAZ0Z |
| chr10 | 4419170 | 4419394 | 98RH.5705C7CE7C5743EZ0Z |
