## Supplementary Table 9 for "An integrated enzymatic and computational pipeline for quantifying off-target base-editing"

**Supplementary Table 9.** **An example of ranking off-target edit sites using *PTPRC* data**

| **weighted score** | **cut cluster** | **chromosome** | **strand** | **cut** | **overlap percentage** | **nearest oncogene** | **nearest oncogene distance** | **nearest gene** | **nearest gene distance** | **in exon** | **in intron** | **levenshtein distance** | **alignment length** | **zscore mean** | **local rank min** | **iguide detect percentage** | **bioinformatic tool detect percentage** |
| --- | --- | --- | --- | --- | --- | --- | --- | --- | --- | --- | --- | --- | --- | --- | --- | --- | --- |
| **51** | **483** | **chr1** | **+** | **198706753** | **96.77** | **PTPRC** | **0** | **PTPRC** | **0** | **1** | **0** | **0** | **20** | **14.03** | **1** | **100** | **85.71** |
| **45** | **6815** | **chr2** | **-** | **143961595** | **100** | **LRP1B** | **1829895** | **GTDC1** | **0** | **0** | **1** | **2** | **20** | **2.92** | **2** | **100** | **100** |
| **41** | **7973** | **chr3** | **+** | **138494327** | **93.55** | **PIK3CB** | **158373** | **CEP70** | **17** | **0** | **0** | **4** | **20** | **3.61** | **2** | **100** | **71.43** |
| **39** | **8490** | **chr3** | **-** | **124196100** | **64.52** | **POLQ** | **2649460** | **KALRN** | **0** | **0** | **1** | **3** | **20** | **0.48** | **4** | **66.67** | **57.14** |
| **37** | **12438** | **chr7** | **+** | **144862723** | **74.19** | **CNTNAP2** | **1253280** | **TPK1** | **26671** | **0** | **0** | **3** | **20** | **0.5** | **4** | **79.17** | **57.14** |
| **36** | **12910** | **chr8** | **+** | **28930553** | **80.65** | **LEPROTL1** | **1164846** | **HMBOX1** | **0** | **0** | **1** | **4** | **20** | **0.67** | **4** | **87.5** | **57.14** |
| **34** | **4224** | **chr15** | **+** | **58902135** | **54.84** | **TCF12** | **1602855** | **SLTM** | **0** | **0** | **1** | **2** | **20** | **0.37** | **2** | **58.33** | **42.86** |
| **29** | **428** | **chr1** | **+** | **183741786** | **51.61** | **TPR** | **2569867** | **RGL1** | **0** | **0** | **1** | **2** | **20** | **1.05** | **5** | **41.67** | **85.71** |
| **26** | **6499** | **chr2** | **-** | **28359446** | **51.61** | **ALK** | **833329** | **BABAM2** | **20546** | **0** | **0** | **3** | **20** | **-0.04** | **6** | **62.5** | **14.29** |
| **25** | **12547** | **chr7** | **-** | **28169228** | **6.45** | **JAZF1** | **0** | **JAZF1** | **0** | **0** | **1** | **2** | **20** | **3.53** | **2** | **0** | **28.57** |
| **25** | **210** | **chr1** | **+** | **88227744** | **22.58** | **BCL10** | **2950655** | **PKN2** | **456529** | **0** | **0** | **3** | **20** | **2.35** | **3** | **12.5** | **57.14** |
| **24** | **1040** | **chr1** | **-** | **204241876** | **12.9** | **MDM4** | **274504** | **PLEKHA6** | **0** | **0** | **1** | **4** | **20** | **2.05** | **4** | **0** | **57.14** |
| **24** | **2923** | **chr12** | **-** | **62039008** | **12.9** | **WIF1** | **3011619** | **TAFA2** | **0** | **0** | **1** | **2** | **21** | **0.56** | **7** | **0** | **57.14** |
| **23** | **3722** | **chr14** | **+** | **40300879** | **45.16** | **FOXA1** | **2704821** | **FBXO33** | **868447** | **0** | **0** | **2** | **20** | **1.24** | **4** | **29.17** | **100** |
| **23** | **9165** | **chr4** | **+** | **174589724** | **29.03** | **CASP3** | **10037973** | **GLRA3** | **47196** | **0** | **0** | **3** | **20** | **1.74** | **7** | **20.83** | **57.14** |
| **23** | **4579** | **chr16** | **+** | **17697707** | **35.48** | **MYH11** | **1840675** | **XYLT1** | **226748** | **0** | **0** | **3** | **20** | **0.74** | **3** | **25** | **71.43** |
| **23** | **5336** | **chr18** | **+** | **57630539** | **38.71** | **MALT1** | **1040848** | **NARS1** | **8704** | **0** | **0** | **3** | **20** | **0.82** | **7** | **29.17** | **71.43** |
| **22** | **12735** | **chr7** | **-** | **115239489** | **16.13** | **MET** | **1432902** | **MDFIC** | **219573** | **0** | **0** | **2** | **20** | **0.78** | **4** | **0** | **71.43** |
| **22** | **6379** | **chr2** | **+** | **218658102** | **38.71** | **FEV** | **322986** | **ZNF142** | **0** | **0** | **1** | **4** | **20** | **0.44** | **6** | **33.33** | **57.14** |
| **21** | **6536** | **chr2** | **-** | **43508376** | **25.81** | **EML4** | **1175829** | **THADA** | **0** | **0** | **1** | **4** | **20** | **0.22** | **8** | **16.67** | **57.14** |
| **20** | **3356** | **chr13** | **+** | **107721970** | **35.48** | **ERCC5** | **4845970** | **NALF1** | **0** | **0** | **1** | **3** | **20** | **0** | **7** | **37.5** | **28.57** |
| **20** | **7645** | **chr3** | **+** | **15096471** | **38.71** | **XPC** | **917689** | **RBSN** | **0** | **0** | **1** | **2** | **18** | **0.19** | **10** | **37.5** | **42.86** |
| **20** | **10033** | **chr4** | **-** | **148700495** | **48.39** | **FBXW7** | **3620050** | **NR3C2** | **255798** | **0** | **0** | **4** | **20** | **0.51** | **8** | **45.83** | **57.14** |
| **20** | **638** | **chr1** | **-** | **22293250** | **9.68** | **ID3** | **1264669** | **WNT4** | **150154** | **0** | **0** | **3** | **20** | **2.41** | **10** | **0** | **42.86** |
| **19** | **8591** | **chr3** | **-** | **164830562** | **16.13** | **MECOM** | **4252938** | **SI** | **148336** | **0** | **0** | **3** | **20** | **1.58** | **8** | **4.17** | **57.14** |
| **19** | **5630** | **chr19** | **+** | **8104809** | **9.68** | **CD209** | **357246** | **FBN3** | **0** | **0** | **1** | **2** | **20** | **0.15** | **10** | **0** | **42.86** |
| **19** | **5654** | **chr19** | **+** | **23485493** | **19.35** | **ZNF429** | **1929224** | **ZNF91** | **90023** | **0** | **0** | **4** | **20** | **0.52** | **4** | **8.33** | **57.14** |
| **19** | **3853** | **chr14** | **+** | **93249161** | **9.68** | **GOLGA5** | **409199** | **BTBD7** | **0** | **0** | **1** | **4** | **20** | **2** | **6** | **0** | **42.86** |
| **18** | **14930** | **chrX** | **-** | **33269841** | **6.45** | **FAM47C** | **3738557** | **DMD** | **0** | **0** | **1** | **4** | **20** | **3.86** | **6** | **0** | **28.57** |
| **18** | **2590** | **chr12** | **+** | **50683821** | **6.45** | **ATF1** | **79890** | **DIP2B** | **0** | **0** | **1** | **4** | **20** | **3.56** | **5** | **0** | **28.57** |
| **18** | **8895** | **chr4** | **+** | **76333768** | **6.45** | **PTPN13** | **10260548** | **CCDC158** | **0** | **0** | **1** | **4** | **20** | **3.24** | **9** | **0** | **28.57** |
| **18** | **7963** | **chr3** | **+** | **133161599** | **6.45** | **STAG1** | **3174635** | **TMEM108** | **0** | **0** | **1** | **4** | **20** | **3.4** | **8** | **0** | **28.57** |
| **18** | **12606** | **chr7** | **-** | **53591737** | **6.45** | **EGFR** | **1427285** | **POM121L12** | **554813** | **0** | **0** | **3** | **20** | **1.62** | **9** | **0** | **28.57** |
| **18** | **200** | **chr1** | **+** | **85158674** | **9.68** | **BCL10** | **107575** | **SYDE2** | **0** | **1** | **0** | **3** | **20** | **1.72** | **16** | **0** | **42.86** |
| **18** | **8510** | **chr3** | **-** | **133903434** | **32.26** | **STAG1** | **2432800** | **RAB6B** | **7553** | **0** | **0** | **3** | **20** | **0.52** | **15** | **25** | **57.14** |
| **18** | **8627** | **chr3** | **-** | **175565663** | **16.13** | **TBL1XR1** | **1453678** | **NAALADL2** | **0** | **0** | **1** | **3** | **20** | **0.05** | **8** | **4.17** | **57.14** |
| **17** | **2632** | **chr12** | **+** | **72915461** | **22.58** | **PTPRB** | **2278022** | **TRHDE** | **244704** | **0** | **0** | **1** | **20** | **0.5** | **3** | **16.67** | **42.86** |
| **17** | **6977** | **chr2** | **-** | **192292150** | **12.9** | **PMS1** | **2414522** | **TMEFF2** | **97218** | **0** | **0** | **3** | **18** | **0.35** | **9** | **8.33** | **28.57** |
| **17** | **13209** | **chr8** | **+** | **131893945** | **22.58** | **NDRG1** | **1343227** | **EFR3A** | **10148** | **0** | **0** | **3** | **20** | **0.75** | **36** | **12.5** | **57.14** |
| **17** | **3517** | **chr13** | **-** | **72141570** | **22.58** | **GPC5** | **19257038** | **DACH1** | **274367** | **0** | **0** | **3** | **20** | **0.62** | **11** | **12.5** | **57.14** |
| **17** | **3933** | **chr14** | **-** | **38950535** | **9.68** | **FOXA1** | **1354477** | **SEC23A** | **81384** | **0** | **0** | **4** | **20** | **2.61** | **7** | **0** | **42.86** |
| **17** | **255** | **chr1** | **+** | **102031561** | **12.9** | **RBM15** | **8306946** | **OLFM3** | **34636** | **0** | **0** | **2** | **18** | **0.59** | **4** | **8.33** | **28.57** |
| **17** | **14459** | **chr9** | **-** | **117461073** | **12.9** | **TNC** | **2342817** | **ASTN2** | **46017** | **0** | **0** | **3** | **20** | **0.21** | **76** | **0** | **57.14** |
| **17** | **4810** | **chr16** | **-** | **61308507** | **16.13** | **CDH11** | **3635247** | **CDH8** | **338743** | **0** | **0** | **2** | **21** | **0.53** | **9** | **8.33** | **42.86** |
| **17** | **13845** | **chr9** | **+** | **112426071** | **6.45** | **TNC** | **2593508** | **HSDL2** | **0** | **0** | **1** | **3** | **20** | **0.77** | **7** | **0** | **28.57** |
| **17** | **15067** | **chrX** | **-** | **89307366** | **9.68** | **ATRX** | **11521098** | **CPXCR1** | **552586** | **0** | **0** | **3** | **20** | **0.66** | **7** | **0** | **42.86** |
| **17** | **15267** | **chrY** | **-** | **6641502** | **9.68** | **not_detected** | **-1** | **AMELY** | **224416** | **0** | **0** | **3** | **20** | **0.66** | **8** | **0** | **42.86** |
| **17** | **13343** | **chr8** | **-** | **49229653** | **9.68** | **TCEA1** | **4736900** | **PPDPFL** | **153561** | **0** | **0** | **3** | **20** | **0.5** | **10** | **0** | **42.86** |
| **16** | **13110** | **chr8** | **+** | **102345566** | **3.23** | **UBR5** | **0** | **UBR5** | **0** | **0** | **1** | **4** | **20** | **2.15** | **6** | **0** | **14.29** |
| **16** | **10221** | **chr5** | **+** | **27172314** | **19.35** | **CDH10** | **2527337** | **CDH9** | **133729** | **0** | **0** | **3** | **20** | **1.21** | **5** | **4.17** | **71.43** |
| **16** | **5477** | **chr18** | **-** | **33998198** | **25.81** | **SS18** | **7906982** | **NOL4** | **0** | **0** | **1** | **4** | **20** | **0.74** | **14** | **12.5** | **71.43** |
| **16** | **6245** | **chr2** | **+** | **172943886** | **12.9** | **HOXD13** | **3149006** | **RAPGEF4** | **0** | **0** | **1** | **3** | **20** | **2.61** | **3** | **4.17** | **42.86** |
| **16** | **3057** | **chr12** | **-** | **112882699** | **19.35** | **PTPN11** | **372787** | **RPH3A** | **0** | **0** | **1** | **4** | **20** | **0.66** | **71** | **8.33** | **57.14** |
| **16** | **15217** | **chrX** | **-** | **154742904** | **19.35** | **MTCP1** | **318719** | **GAB3** | **0** | **0** | **1** | **4** | **20** | **0.03** | **95** | **8.33** | **57.14** |
| **15** | **5591** | **chr18** | **-** | **72164935** | **9.68** | **KDSR** | **8797426** | **CBLN2** | **371746** | **0** | **0** | **3** | **20** | **2.08** | **18** | **0** | **42.86** |
| **15** | **7145** | **chr20** | **+** | **11019572** | **6.45** | **CRNKL1** | **9014797** | **JAG1** | **345574** | **0** | **0** | **5** | **20** | **1.86** | **9** | **0** | **28.57** |
| **15** | **13062** | **chr8** | **+** | **83375226** | **6.45** | **CNBD1** | **3491217** | **RALYL** | **807561** | **0** | **0** | **2** | **20** | **0.67** | **8** | **0** | **28.57** |
| **15** | **6951** | **chr2** | **-** | **184581393** | **6.45** | **ITGAV** | **2008673** | **ZNF804A** | **17136** | **0** | **0** | **4** | **20** | **3.06** | **5** | **0** | **28.57** |
| **15** | **6162** | **chr2** | **+** | **141740395** | **9.68** | **LRP1B** | **0** | **LRP1B** | **0** | **0** | **1** | **4** | **20** | **0.69** | **84** | **0** | **42.86** |
| **15** | **159** | **chr1** | **+** | **72912278** | **6.45** | **FUBP1** | **5031778** | **NEGR1** | **629740** | **0** | **0** | **4** | **20** | **3.89** | **1** | **0** | **28.57** |
| **15** | **5201** | **chr17** | **-** | **79787761** | **9.68** | **RNF213** | **473106** | **CBX2** | **0** | **1** | **0** | **3** | **20** | **0.26** | **29** | **0** | **42.86** |
| **15** | **13513** | **chr8** | **-** | **113179145** | **9.68** | **CSMD3** | **0** | **CSMD3** | **0** | **0** | **1** | **4** | **20** | **-0.3** | **132** | **0** | **42.86** |
| **15** | **8885** | **chr4** | **+** | **73738512** | **6.45** | **PTPN13** | **12855804** | **CXCL8** | **2057** | **0** | **0** | **3** | **20** | **0.74** | **8** | **0** | **28.57** |
| **14** | **11371** | **chr6** | **+** | **105097903** | **6.45** | **PRDM1** | **988418** | **BVES** | **0** | **1** | **0** | **4** | **20** | **2.21** | **26** | **0** | **28.57** |
| **14** | **10970** | **chr5** | **-** | **124074003** | **25.81** | **ACSL6** | **7875971** | **CSNK1G3** | **456955** | **0** | **0** | **4** | **20** | **1.02** | **15** | **16.67** | **57.14** |
| **14** | **4189** | **chr15** | **+** | **49245406** | **29.03** | **USP8** | **1178975** | **GALK2** | **0** | **0** | **1** | **4** | **20** | **0.32** | **11** | **20.83** | **57.14** |
| **14** | **1999** | **chr11** | **+** | **78443451** | **9.68** | **NUMA1** | **6362759** | **NARS2** | **0** | **0** | **1** | **3** | **20** | **1.07** | **46** | **0** | **42.86** |
| **14** | **12012** | **chr6** | **-** | **166707907** | **6.45** | **FGFR1OP** | **291276** | **RPS6KA2** | **0** | **0** | **1** | **4** | **20** | **-0.06** | **9** | **8.33** | **0** |
| **14** | **5229** | **chr18** | **+** | **7811120** | **9.68** | **ZNF521** | **17250807** | **PTPRM** | **0** | **0** | **1** | **3** | **20** | **0.91** | **23** | **0** | **42.86** |
| **14** | **8598** | **chr3** | **-** | **168109846** | **12.9** | **MECOM** | **973654** | **GOLIM4** | **13923** | **0** | **0** | **4** | **20** | **1.02** | **15** | **0** | **57.14** |
| **14** | **12450** | **chr7** | **+** | **147998717** | **3.23** | **CNTNAP2** | **0** | **CNTNAP2** | **0** | **0** | **1** | **2** | **21** | **1.97** | **148** | **0** | **14.29** |
| **14** | **14366** | **chr9** | **-** | **81378935** | **12.9** | **NTRK2** | **3289617** | **TLE1** | **204748** | **0** | **0** | **4** | **20** | **0.32** | **88** | **0** | **57.14** |
| **14** | **133** | **chr1** | **+** | **66705466** | **9.68** | **JAK1** | **1738963** | **SGIP1** | **0** | **0** | **1** | **3** | **20** | **0.01** | **152** | **0** | **42.86** |
| **14** | **13763** | **chr9** | **+** | **74628516** | **9.68** | **GNAQ** | **3087572** | **RORB** | **0** | **0** | **1** | **3** | **20** | **-0.05** | **90** | **0** | **42.86** |
| **14** | **10392** | **chr5** | **+** | **81289788** | **22.58** | **RAD17** | **11874988** | **ZCCHC9** | **11799** | **0** | **0** | **4** | **20** | **0.07** | **149** | **12.5** | **57.14** |
| **14** | **11514** | **chr6** | **+** | **157035211** | **3.23** | **ARID1B** | **0** | **ARID1B** | **0** | **0** | **1** | **3** | **20** | **1.58** | **256** | **0** | **14.29** |
| **14** | **1302** | **chr10** | **+** | **50570294** | **16.13** | **A1CF** | **229116** | **SGMS1** | **0** | **0** | **1** | **2** | **20** | **-0.07** | **39** | **20.83** | **0** |
| **13** | **6939** | **chr2** | **-** | **181846701** | **3.23** | **NFE2L2** | **4454005** | **ITPRID2** | **45029** | **0** | **0** | **3** | **21** | **7.62** | **4** | **0** | **14.29** |
| **13** | **7610** | **chr21** | **-** | **44527735** | **6.45** | **U2AF1** | **1420149** | **TSPEAR** | **0** | **0** | **1** | **4** | **20** | **2.42** | **27** | **0** | **28.57** |
| **13** | **10237** | **chr5** | **+** | **32051408** | **6.45** | **GOLPH3** | **73297** | **PDZD2** | **0** | **0** | **1** | **4** | **20** | **2.44** | **24** | **0** | **28.57** |
| **13** | **4174** | **chr15** | **+** | **47646958** | **6.45** | **HMGN2P46** | **2060669** | **SEMA6D** | **0** | **0** | **1** | **4** | **20** | **2.29** | **25** | **0** | **28.57** |
| **13** | **1101** | **chr1** | **-** | **222931281** | **6.45** | **H3F3A** | **3130571** | **DISP1** | **0** | **0** | **1** | **4** | **20** | **2.11** | **28** | **0** | **28.57** |
| **13** | **1495** | **chr10** | **-** | **289266** | **6.45** | **LARP4B** | **517649** | **DIP2C** | **0** | **0** | **1** | **4** | **20** | **2.11** | **30** | **0** | **28.57** |
| **13** | **6345** | **chr2** | **+** | **203121673** | **6.45** | **CD28** | **584803** | **NBEAL1** | **0** | **0** | **1** | **4** | **20** | **2.11** | **29** | **0** | **28.57** |
| **13** | **9940** | **chr4** | **-** | **115821523** | **32.26** | **IL2** | **6629948** | **NDST4** | **707904** | **0** | **0** | **3** | **18** | **-0.06** | **15** | **29.17** | **42.86** |
| **13** | **11194** | **chr6** | **+** | **41983875** | **6.45** | **CCND3** | **0** | **CCND3** | **0** | **0** | **1** | **5** | **20** | **-0.11** | **58** | **8.33** | **0** |
| **13** | **11627** | **chr6** | **-** | **34598774** | **12.9** | **HMGA1** | **352544** | **ILRUN** | **0** | **0** | **1** | **4** | **20** | **2.16** | **7** | **4.17** | **42.86** |
| **13** | **6479** | **chr2** | **-** | **21561763** | **6.45** | **WDCP** | **2467578** | **TDRD15** | **417396** | **0** | **0** | **3** | **20** | **1.57** | **56** | **0** | **28.57** |
| **13** | **2361** | **chr11** | **-** | **96334118** | **6.45** | **MAML2** | **0** | **MAML2** | **0** | **0** | **1** | **6** | **20** | **-0.07** | **33** | **8.33** | **0** |
| **13** | **3710** | **chr14** | **+** | **37589748** | **3.23** | **FOXA1** | **237** | **FOXA1** | **0** | **1** | **0** | **3** | **20** | **1.66** | **231** | **0** | **14.29** |
| **13** | **12290** | **chr7** | **+** | **99049423** | **6.45** | **TRRAP** | **0** | **SMURF1** | **0** | **0** | **1** | **6** | **20** | **-0.04** | **124** | **8.33** | **0** |
| **13** | **2758** | **chr12** | **+** | **122362301** | **6.45** | **CLIP1** | **0** | **CLIP1** | **0** | **0** | **1** | **6** | **20** | **-0.03** | **21** | **8.33** | **0** |
| **13** | **1839** | **chr11** | **+** | **3744854** | **6.45** | **NUP98** | **0** | **NUP98** | **0** | **0** | **1** | **6** | **20** | **-0.01** | **181** | **8.33** | **0** |
| **13** | **1244** | **chr10** | **+** | **26786651** | **6.45** | **ABI1** | **0** | **ABI1** | **0** | **0** | **1** | **6** | **19** | **-0.06** | **358** | **8.33** | **0** |
| **13** | **9104** | **chr4** | **+** | **152451637** | **6.45** | **FBXW7** | **0** | **FBXW7** | **0** | **0** | **1** | **4** | **20** | **-0.21** | **211** | **0** | **28.57** |
| **13** | **12948** | **chr8** | **+** | **42982497** | **6.45** | **HOOK3** | **0** | **HOOK3** | **0** | **0** | **1** | **6** | **20** | **-0.06** | **447** | **8.33** | **0** |
| **13** | **13613** | **chr9** | **+** | **10210360** | **6.45** | **PTPRD** | **0** | **PTPRD** | **0** | **0** | **1** | **6** | **20** | **-0.06** | **441** | **8.33** | **0** |
| **13** | **8923** | **chr4** | **+** | **87080920** | **6.45** | **AFF1** | **0** | **AFF1** | **0** | **0** | **1** | **6** | **20** | **-0.04** | **296** | **8.33** | **0** |
| **13** | **3179** | **chr13** | **+** | **46138584** | **6.45** | **LCP1** | **0** | **LCP1** | **0** | **0** | **1** | **6** | **20** | **-0.02** | **47** | **8.33** | **0** |
| **12** | **7786** | **chr3** | **+** | **65866815** | **3.23** | **MITF** | **3872621** | **MAGI1** | **0** | **0** | **1** | **6** | **21** | **6.48** | **2** | **0** | **14.29** |
| **12** | **12506** | **chr7** | **-** | **14803347** | **3.23** | **ETV1** | **811923** | **DGKB** | **0** | **0** | **1** | **5** | **20** | **3.7** | **10** | **0** | **14.29** |
| **12** | **3374** | **chr13** | **-** | **21942310** | **9.68** | **LATS2** | **880764** | **FGF9** | **237813** | **0** | **0** | **3** | **21** | **0.88** | **18** | **0** | **42.86** |
| **12** | **7060** | **chr2** | **-** | **220674332** | **9.68** | **PAX3** | **1525557** | **EPHA4** | **743695** | **0** | **0** | **3** | **21** | **1.18** | **21** | **0** | **42.86** |
| **12** | **3809** | **chr14** | **+** | **75894855** | **6.45** | **TSHR** | **5060135** | **TTLL5** | **0** | **0** | **1** | **3** | **21** | **0.13** | **36** | **0** | **28.57** |
| **12** | **14408** | **chr9** | **-** | **95364904** | **3.23** | **FANCC** | **0** | **FANCC** | **47196** | **0** | **0** | **3** | **19** | **2.78** | **67** | **0** | **14.29** |
| **12** | **10181** | **chr5** | **+** | **10372151** | **6.45** | **CTNND2** | **599690** | **MARCHF6** | **0** | **0** | **1** | **4** | **20** | **1.51** | **59** | **0** | **28.57** |
| **12** | **2434** | **chr11** | **-** | **122210267** | **9.68** | **ARHGEF12** | **1720332** | **BLID** | **94053** | **0** | **0** | **3** | **20** | **0.65** | **51** | **0** | **42.86** |
| **12** | **5819** | **chr19** | **-** | **57248978** | **32.26** | **CNOT3** | **3093271** | **ZNF805** | **0** | **0** | **1** | **5** | **20** | **-0.02** | **26** | **33.33** | **28.57** |
| **12** | **7125** | **chr20** | **+** | **3031933** | **6.45** | **SIRPA** | **1091342** | **PTPRA** | **0** | **0** | **1** | **3** | **19** | **-0.07** | **75** | **8.33** | **0** |
| **12** | **4049** | **chr14** | **-** | **76697756** | **9.68** | **TSHR** | **4257234** | **VASH1** | **63712** | **0** | **0** | **3** | **20** | **0.43** | **54** | **0** | **42.86** |
| **12** | **12858** | **chr8** | **+** | **8749214** | **9.68** | **ARHGEF10** | **6790574** | **MFHAS1** | **34140** | **0** | **0** | **3** | **20** | **0.66** | **27** | **0** | **42.86** |
| **12** | **835** | **chr1** | **-** | **104194320** | **9.68** | **RBM15** | **6144187** | **AMY1C** | **435629** | **0** | **0** | **3** | **20** | **0.21** | **53** | **0** | **42.86** |
| **12** | **8197** | **chr3** | **-** | **23158264** | **9.68** | **TGFBR2** | **7448239** | **UBE2E2** | **44834** | **0** | **0** | **3** | **20** | **0.16** | **81** | **0** | **42.86** |
| **12** | **4202** | **chr15** | **+** | **52564774** | **6.45** | **MYO5A** | **35725** | **ARPP19** | **0** | **0** | **1** | **3** | **20** | **-0.61** | **84** | **0** | **28.57** |
| **12** | **244** | **chr1** | **+** | **97865367** | **6.45** | **RPL5** | **5023444** | **DPYD** | **0** | **0** | **1** | **3** | **20** | **0.39** | **34** | **0** | **28.57** |
| **12** | **7942** | **chr3** | **+** | **126650493** | **6.45** | **GATA2** | **1828935** | **TXNRD3** | **0** | **0** | **1** | **3** | **20** | **-0.16** | **87** | **0** | **28.57** |
| **12** | **8189** | **chr3** | **-** | **19198728** | **6.45** | **XPC** | **5019946** | **KCNH8** | **0** | **0** | **1** | **3** | **20** | **-0.13** | **161** | **0** | **28.57** |
| **12** | **12559** | **chr7** | **-** | **34806769** | **6.45** | **FKBP9** | **1799839** | **NPSR1** | **0** | **0** | **1** | **3** | **20** | **-0.19** | **103** | **0** | **28.57** |
| **12** | **8621** | **chr3** | **-** | **173940751** | **6.45** | **TBL1XR1** | **3078590** | **NLGN1** | **0** | **0** | **1** | **3** | **20** | **0.9** | **17** | **0** | **28.57** |
| **12** | **759** | **chr1** | **-** | **76548797** | **6.45** | **FUBP1** | **1395259** | **ST6GALNAC3** | **0** | **0** | **1** | **3** | **20** | **-0.15** | **56** | **0** | **28.57** |
| **12** | **1881** | **chr11** | **+** | **24713125** | **6.45** | **FANCF** | **2086339** | **LUZP2** | **0** | **0** | **1** | **3** | **20** | **-0.35** | **175** | **0** | **28.57** |
| **12** | **2808** | **chr12** | **-** | **13857255** | **6.45** | **CDKN1B** | **1134885** | **GRIN2B** | **0** | **0** | **1** | **3** | **20** | **-0.13** | **86** | **0** | **28.57** |
| **12** | **2946** | **chr12** | **-** | **70318598** | **6.45** | **PTPRB** | **197269** | **CNOT2** | **0** | **0** | **1** | **3** | **18** | **0.58** | **29** | **0** | **28.57** |
| **12** | **849** | **chr1** | **-** | **107262083** | **6.45** | **RBM15** | **3076424** | **NTNG1** | **0** | **0** | **1** | **3** | **20** | **-0.37** | **228** | **0** | **28.57** |
| **12** | **3180** | **chr13** | **+** | **47423665** | **6.45** | **RB1** | **880062** | **SUCLA2** | **518991** | **0** | **0** | **6** | **19** | **-0.13** | **8** | **8.33** | **0** |
| **12** | **4726** | **chr16** | **-** | **7075191** | **6.45** | **GRIN2A** | **2678214** | **RBFOX1** | **0** | **0** | **1** | **3** | **20** | **0.34** | **31** | **0** | **28.57** |
| **12** | **6603** | **chr2** | **-** | **64529397** | **6.45** | **XPO1** | **2990772** | **AFTPH** | **0** | **0** | **1** | **3** | **20** | **0.12** | **49** | **0** | **28.57** |
| **12** | **4054** | **chr14** | **-** | **79004825** | **6.45** | **TSHR** | **1950165** | **NRXN3** | **0** | **0** | **1** | **3** | **20** | **0.07** | **54** | **0** | **28.57** |
| **12** | **722** | **chr1** | **-** | **64967930** | **6.45** | **JAK1** | **1427** | **JAK1** | **0** | **0** | **1** | **3** | **21** | **-0.49** | **256** | **0** | **28.57** |
| **12** | **5627** | **chr19** | **+** | **6749412** | **6.45** | **VAV1** | **23303** | **TRIP10** | **0** | **0** | **1** | **3** | **20** | **-0.48** | **345** | **0** | **28.57** |
| **12** | **11762** | **chr6** | **-** | **83895849** | **6.45** | **EPHA7** | **9344172** | **RIPPLY2-CYB5R4** | **0** | **0** | **1** | **3** | **20** | **-0.45** | **79** | **0** | **28.57** |
| **12** | **1802** | **chr10** | **-** | **115220608** | **6.45** | **SHTN1** | **1660875** | **ATRNL1** | **0** | **0** | **1** | **3** | **18** | **0.19** | **18** | **0** | **28.57** |
| **12** | **8270** | **chr3** | **-** | **50688876** | **6.45** | **RHOA** | **1275879** | **DOCK3** | **0** | **0** | **1** | **3** | **18** | **0.19** | **22** | **0** | **28.57** |
| **12** | **1240** | **chr10** | **+** | **24622785** | **6.45** | **ABI1** | **2123809** | **ARHGAP21** | **0** | **1** | **1** | **6** | **20** | **-0.04** | **669** | **8.33** | **0** |
| **11** | **6728** | **chr2** | **-** | **109185311** | **3.23** | **RANBP2** | **399501** | **SH3RF3** | **0** | **0** | **1** | **1** | **20** | **0.99** | **2** | **0** | **14.29** |
| **11** | **14359** | **chr9** | **-** | **78885084** | **16.13** | **GNAQ** | **853627** | **PSAT1** | **554992** | **0** | **0** | **2** | **19** | **0.72** | **4** | **16.67** | **14.29** |
| **11** | **4019** | **chr14** | **-** | **67833478** | **3.23** | **RAD51B** | **0** | **RAD51B** | **0** | **0** | **1** | **4** | **21** | **4.03** | **13** | **0** | **14.29** |
| **11** | **5202** | **chr17** | **-** | **79787761** | **3.23** | **RNF213** | **473106** | **CBX2** | **0** | **1** | **0** | **3** | **20** | **3.29** | **24** | **0** | **14.29** |
| **11** | **5563** | **chr18** | **-** | **65517581** | **6.45** | **KDSR** | **2150072** | **CDH7** | **232671** | **0** | **0** | **4** | **20** | **2.75** | **19** | **0** | **28.57** |
| **11** | **13264** | **chr8** | **-** | **11957047** | **6.45** | **PCM1** | **5965794** | **DEFB136** | **16890** | **0** | **0** | **4** | **20** | **2.59** | **20** | **0** | **28.57** |
| **11** | **7139** | **chr20** | **+** | **10198474** | **6.45** | **SIRPA** | **8257883** | **SNAP25** | **20356** | **0** | **0** | **4** | **20** | **2.59** | **21** | **0** | **28.57** |
| **11** | **13625** | **chr9** | **+** | **14294329** | **3.23** | **NFIB** | **0** | **NFIB** | **0** | **0** | **1** | **4** | **22** | **3.12** | **39** | **0** | **14.29** |
| **11** | **4975** | **chr17** | **+** | **55279637** | **3.23** | **HLF** | **0** | **HLF** | **0** | **0** | **1** | **5** | **20** | **2.09** | **48** | **0** | **14.29** |
| **11** | **2468** | **chr12** | **+** | **4690779** | **3.23** | **CCND2** | **385430** | **NDUFA9** | **0** | **1** | **0** | **3** | **20** | **2.95** | **50** | **0** | **14.29** |
| **11** | **13215** | **chr8** | **+** | **133259524** | **3.23** | **NDRG1** | **0** | **NDRG1** | **0** | **0** | **1** | **4** | **20** | **1.92** | **55** | **0** | **14.29** |
| **11** | **7170** | **chr20** | **+** | **18374746** | **6.45** | **CRNKL1** | **1659623** | **DZANK1** | **8621** | **0** | **0** | **4** | **20** | **1.9** | **39** | **0** | **28.57** |
| **11** | **201** | **chr1** | **+** | **85158674** | **3.23** | **BCL10** | **107575** | **SYDE2** | **0** | **1** | **0** | **3** | **20** | **2.86** | **58** | **0** | **14.29** |
| **11** | **11914** | **chr6** | **-** | **131893145** | **3.23** | **SGK1** | **2276102** | **ENPP1** | **0** | **1** | **0** | **3** | **20** | **2.59** | **82** | **0** | **14.29** |
| **11** | **11847** | **chr6** | **-** | **111580520** | **12.9** | **FOXO3** | **2895747** | **TRAF3IP2** | **0** | **0** | **1** | **4** | **21** | **-0.28** | **42** | **8.33** | **28.57** |
| **11** | **5449** | **chr18** | **-** | **24090392** | **9.68** | **ZNF521** | **971535** | **TTC39C** | **0** | **0** | **1** | **4** | **20** | **0.99** | **54** | **0** | **42.86** |
