## Supplementary Table 10 for "An integrated enzymatic and computational pipeline for quantifying off-target base-editing"

**Supplementary Table 10.** **An example of choices of weights for ranking off-target sites**

| **variable** | **variable_type** | **source** | **type** | **condition** | **lower_threshold** | **upper_threshold** | **weight** |
| --- | --- | --- | --- | --- | --- | --- | --- |
| **overlap_percentage** | **overlap** | **default** | **feature** | **>=** | **90** | **100** | **10** |
| **overlap_percentage** | **overlap** | **default** | **feature** | **>=** | **50** | **89** | **7** |
| **overlap_percentage** | **overlap** | **default** | **feature** | **>=** | **30** | **49** | **3** |
| **overlap_percentage** | **overlap** | **default** | **feature** | **>=** | **7** | **29** | **2** |
| **zscore_mean** | **presence** | **default** | **feature** | **>=** | **1.5** | **inf** | **3** |
| **zscore_min** | **presence** | **default** | **feature** | **>=** | **1.5** | **inf** | **1** |
| **local_rank_min** | **presence** | **default** | **feature** | **>=** | **0** | **10** | **5** |
| **levenshtein_distance** | **distance** | **default** | **feature** | **>=** | **0** | **3** | **8** |
| **levenshtein_distance** | **distance** | **default** | **feature** | **>=** | **4** | **6** | **5** |
| **alignment_length** | **distance** | **default** | **feature** | **>=** | **17** | **22** | **2** |
| **bioinformatic_tool_detect_percentage** | **overlap** | **user** | **method** | **>=** | **50** | **100** | **5** |
| **bioinformatic_tool_detect_percentage** | **overlap** | **user** | **method** | **>=** | **1** | **15** | **-6** |
| **iguide_detect_percentage** | **overlap** | **user** | **method** | **>=** | **1** | **5** | **-6** |
| **iguide_detect_percentage** | **overlap** | **user** | **method** | **>=** | **50** | **100** | **10** |
| **nearest_gene_distance** | **distance** | **user** | **feature** | **>=** | **1** | **1000** | **1** |
| **in_exon** | **presence** | **user** | **feature** | **==** | **1** | **inf** | **3** |
| **in_intron** | **presence** | **user** | **feature** | **==** | **1** | **inf** | **2** |
| **nearest_oncogene_distance** | **distance** | **user** | **feature** | **==** | **0** | **0** | **4** |
| **nearest_oncogene_distance** | **distance** | **user** | **feature** | **>=** | **1** | **1000** | **2** |
