## Supplementary Table 11 for "An integrated enzymatic and computational pipeline for quantifying off-target base-editing"

**Supplementary Table 11.** **Oligonucleotides**

| **Oligonucleotide Name** | **Sequence** | **Company** | **Reference** |
| --- | --- | --- | --- |
| sgPTPRC - sgRNA protospacer sequence | 5’- AAAATATGCAAACATCACTG -3’ | Integrated DNA Technologies (IDT) or Synthego, standard desalting | PMID: 37651540 |
| sgPTPRC PCR Primer Forward, | 5’- ACAAGCTGAGGTCCTTGTTAGG -3’ | Integrated DNA Technologies (IDT), standard desalting | PMID: 37651540 |
| sgPTPRC PCR Primer Reverse, | 5’- AGCTTGCAGACAATCACTTAGC -3’ | Integrated DNA Technologies (IDT), standard desalting | PMID: 37651540 |
| sgPTPRC Sequence Primer Forward, | 5’- GGCAGTAAAGCACAGATAATTACG -3’ | Integrated DNA Technologies (IDT), standard desalting | PMID: 37651540 |
| **dsODN oligos** |  |  |  |
| dsODN Oligonucleotide Sense | /5Phos/G*C*TCGCGTTTAATTGAGTTGTCATATGTTAATAACGGTATACGC*G*A | Integrated DNA Technologies (IDT), Duplex with HPLC Purification | PMID: 30654827 |
| dsODN Oligonucleotide Antisense | /5Phos/T*C*GCGTATACCGTTATTAACATATGACAACTCAATTAAACGCGA*G*C | Integrated DNA Technologies (IDT), Duplex with HPLC Purification | PMID: 30654827 |
| **Adaptor oligos** |  |  |  |
| adapter_common_oligo | /5Phos/GATCGGAAGAGC*C*A | Integrated DNA Technologies (IDT), standard desalting | PMID: 25513782 |
| A01 | AATGATACGGCGACCACCGAGATCTACACTAGATCGCNNWNNWNNACACTCTTTCCCTACACGACGCTCTTCCGATC*T | Integrated DNA Technologies (IDT), standard desalting | PMID: 25513782 |
| A02 | AATGATACGGCGACCACCGAGATCTACACCTCTCTATNNWNNWNNACACTCTTTCCCTACACGACGCTCTTCCGATC*T | Integrated DNA Technologies (IDT), standard desalting | PMID: 25513782 |
| A03 | AATGATACGGCGACCACCGAGATCTACACTATCCTCTNNWNNWNNACACTCTTTCCCTACACGACGCTCTTCCGATC*T | Integrated DNA Technologies (IDT), standard desalting | PMID: 25513782 |
| A04 | AATGATACGGCGACCACCGAGATCTACACAGAGTAGANNWNNWNNACACTCTTTCCCTACACGACGCTCTTCCGATC*T | Integrated DNA Technologies (IDT), standard desalting | PMID: 25513782 |
| A05 | AATGATACGGCGACCACCGAGATCTACACGTAAGGAGNNWNNWNNACACTCTTTCCCTACACGACGCTCTTCCGATC*T | Integrated DNA Technologies (IDT), standard desalting | PMID: 25513782 |
| A06 | AATGATACGGCGACCACCGAGATCTACACACTGCATANNWNNWNNACACTCTTTCCCTACACGACGCTCTTCCGATC*T | Integrated DNA Technologies (IDT), standard desalting | PMID: 25513782 |
| A07 | AATGATACGGCGACCACCGAGATCTACACAAGGAGTANNWNNWNNACACTCTTTCCCTACACGACGCTCTTCCGATC*T | Integrated DNA Technologies (IDT), standard desalting | PMID: 25513782 |
| A08 | AATGATACGGCGACCACCGAGATCTACACCTAAGCCTNNWNNWNNACACTCTTTCCCTACACGACGCTCTTCCGATC*T | Integrated DNA Technologies (IDT), standard desalting | PMID: 25513782 |
| A09 | AATGATACGGCGACCACCGAGATCTACACGACATTGTNNWNNWNNACACTCTTTCCCTACACGACGCTCTTCCGATC*T | Integrated DNA Technologies (IDT), standard desalting | PMID: 25513782 |
| A10 | AATGATACGGCGACCACCGAGATCTACACACTGATGGNNWNNWNNACACTCTTTCCCTACACGACGCTCTTCCGATC*T | Integrated DNA Technologies (IDT), standard desalting | PMID: 25513782 |
| A11 | AATGATACGGCGACCACCGAGATCTACACGTACCTAGNNWNNWNNACACTCTTTCCCTACACGACGCTCTTCCGATC*T | Integrated DNA Technologies (IDT), standard desalting | PMID: 25513782 |
| A12 | AATGATACGGCGACCACCGAGATCTACACCAGAGCTANNWNNWNNACACTCTTTCCCTACACGACGCTCTTCCGATC*T | Integrated DNA Technologies (IDT), standard desalting | PMID: 25513782 |
| A13 | AATGATACGGCGACCACCGAGATCTACACCATAGTGANNWNNWNNACACTCTTTCCCTACACGACGCTCTTCCGATC*T | Integrated DNA Technologies (IDT), standard desalting | PMID: 25513782 |
| A14 | AATGATACGGCGACCACCGAGATCTACACTACCTAGTNNWNNWNNACACTCTTTCCCTACACGACGCTCTTCCGATC*T | Integrated DNA Technologies (IDT), standard desalting | PMID: 25513782 |
| A15 | AATGATACGGCGACCACCGAGATCTACACCGCGATATNNWNNWNNACACTCTTTCCCTACACGACGCTCTTCCGATC*T | Integrated DNA Technologies (IDT), standard desalting | PMID: 25513782 |
| A16 | AATGATACGGCGACCACCGAGATCTACACTGGATTGTNNWNNWNNACACTCTTTCCCTACACGACGCTCTTCCGATC*T | Integrated DNA Technologies (IDT), standard desalting | PMID: 25513782 |
| **Adapter Primers** |  |  |  |
| P5_1 | AATGATACGGCGACCACCGAGATCTA | Integrated DNA Technologies (IDT), standard desalting | PMID: 25513782 |
| P5_2 | AATGATACGGCGACCACCGAGATCTACAC | Integrated DNA Technologies (IDT), standard desalting | PMID: 25513782 |
| **dsODN Primers** |  |  |  |
| GSP1_pos_Nuc_off | GGATCTCGACGCTCTCCCTATACCGTTATTAACATATGACA | Integrated DNA Technologies (IDT), standard desalting | PMID: 25513782 |
| GSP1_neg_Nuc_off | GGATCTCGACGCTCTCCCTGTTTAATTGAGTTGTCATATGTTAATAAC | Integrated DNA Technologies (IDT), standard desalting | PMID: 25513782 |
| GSP2_pos_Nuc_off | CCTCTCTATGGGCAGTCGGTGATACATATGACAACTCAATTAAAC | Integrated DNA Technologies (IDT), standard desalting | PMID: 25513782 |
| GSP2_neg_Nuc_off | CCTCTCTATGGGCAGTCGGTGATTTGAGTTGTCATATGTTAATAACGGTA | Integrated DNA Technologies (IDT), standard desalting | PMID: 25513782 |
| **Barcoded dsODN Tiered Primers** |  |  |  |
| P7_iGuide_PCR2_701 | CAAGCAGAAGACGGCATACGAGATAGGGCTATAGTTGTGACTGGAGTCCTCTCTATGGGCAGTCGGTGA | Integrated DNA Technologies (IDT), standard desalting | This study |
| P7_iGuide_PCR2_702 | CAAGCAGAAGACGGCATACGAGATTGTCTCGCAAGCGTGACTGGAGTCCTCTCTATGGGCAGTCGGTGA | Integrated DNA Technologies (IDT), standard desalting | This study |
| P7_iGuide_PCR2_703 | CAAGCAGAAGACGGCATACGAGATCAGCCGCATATCGTGACTGGAGTCCTCTCTATGGGCAGTCGGTGA | Integrated DNA Technologies (IDT), standard desalting | This study |
| P7_iGuide_PCR2_704 | CAAGCAGAAGACGGCATACGAGATGATACGTTCGCAGTGACTGGAGTCCTCTCTATGGGCAGTCGGTGA | Integrated DNA Technologies (IDT), standard desalting | This study |
| P7_iGuide_PCR2_705 | CAAGCAGAAGACGGCATACGAGATCCAAGATTCGCCGTGACTGGAGTCCTCTCTATGGGCAGTCGGTGA | Integrated DNA Technologies (IDT), standard desalting | This study |
| P7_iGuide_PCR2_706 | CAAGCAGAAGACGGCATACGAGATGAGGCTGATTTAGTGACTGGAGTCCTCTCTATGGGCAGTCGGTGA | Integrated DNA Technologies (IDT), standard desalting | This study |
| P7_iGuide_PCR2_707 | CAAGCAGAAGACGGCATACGAGATGAGTTAGCATCAGTGACTGGAGTCCTCTCTATGGGCAGTCGGTGA | Integrated DNA Technologies (IDT), standard desalting | This study |
| P7_iGuide_PCR2_708 | CAAGCAGAAGACGGCATACGAGATTGTAGTATAGGCGTGACTGGAGTCCTCTCTATGGGCAGTCGGTGA | Integrated DNA Technologies (IDT), standard desalting | This study |
| P7_iGuide_PCR2_709 | CAAGCAGAAGACGGCATACGAGATCTCACGCAATGCGTGACTGGAGTCCTCTCTATGGGCAGTCGGTGA | Integrated DNA Technologies (IDT), standard desalting | This study |
| P7_iGuide_PCR2_710 | CAAGCAGAAGACGGCATACGAGATGTCCCGTGAAATGTGACTGGAGTCCTCTCTATGGGCAGTCGGTGA | Integrated DNA Technologies (IDT), standard desalting | This study |
| P7_iGuide_PCR2_711 | CAAGCAGAAGACGGCATACGAGATGGACAGTGTATTGTGACTGGAGTCCTCTCTATGGGCAGTCGGTGA | Integrated DNA Technologies (IDT), standard desalting | This study |
| P7_iGuide_PCR2_712 | CAAGCAGAAGACGGCATACGAGATACACGACTATAGGTGACTGGAGTCCTCTCTATGGGCAGTCGGTGA | Integrated DNA Technologies (IDT), standard desalting | This study |
| P7_iGuide_PCR2_713 | CAAGCAGAAGACGGCATACGAGATGTGTAGGTGCTTGTGACTGGAGTCCTCTCTATGGGCAGTCGGTGA | Integrated DNA Technologies (IDT), standard desalting | This study |
| P7_iGuide_PCR2_714 | CAAGCAGAAGACGGCATACGAGATTGAACTAGCGTCGTGACTGGAGTCCTCTCTATGGGCAGTCGGTGA | Integrated DNA Technologies (IDT), standard desalting | This study |
| P7_iGuide_PCR2_715 | CAAGCAGAAGACGGCATACGAGATTCCGAGTCACCAGTGACTGGAGTCCTCTCTATGGGCAGTCGGTGA | Integrated DNA Technologies (IDT), standard desalting | This study |
| P7_iGuide_PCR2_716 | CAAGCAGAAGACGGCATACGAGATTCCTCTTTGGTCGTGACTGGAGTCCTCTCTATGGGCAGTCGGTGA | Integrated DNA Technologies (IDT), standard desalting | This study |
| P7_iGuide_PCR2_717 | CAAGCAGAAGACGGCATACGAGATTCCACCCTCTATGTGACTGGAGTCCTCTCTATGGGCAGTCGGTGA | Integrated DNA Technologies (IDT), standard desalting | This study |
| P7_iGuide_PCR2_718 | CAAGCAGAAGACGGCATACGAGATTCGTGACGCTAAGTGACTGGAGTCCTCTCTATGGGCAGTCGGTGA | Integrated DNA Technologies (IDT), standard desalting | This study |
| P7_iGuide_PCR2_719 | CAAGCAGAAGACGGCATACGAGATACGGCTAGTTCCGTGACTGGAGTCCTCTCTATGGGCAGTCGGTGA | Integrated DNA Technologies (IDT), standard desalting | This study |
| P7_iGuide_PCR2_720 | CAAGCAGAAGACGGCATACGAGATGCACTGGCATATGTGACTGGAGTCCTCTCTATGGGCAGTCGGTGA | Integrated DNA Technologies (IDT), standard desalting | This study |
| P7_iGuide_PCR2_721 | CAAGCAGAAGACGGCATACGAGATGGCATTAGTTGAGTGACTGGAGTCCTCTCTATGGGCAGTCGGTGA | Integrated DNA Technologies (IDT), standard desalting | This study |
| P7_iGuide_PCR2_722 | CAAGCAGAAGACGGCATACGAGATCGGTAGTTGATCGTGACTGGAGTCCTCTCTATGGGCAGTCGGTGA | Integrated DNA Technologies (IDT), standard desalting | This study |
| P7_iGuide_PCR2_723 | CAAGCAGAAGACGGCATACGAGATTGAAAGCGGCGAGTGACTGGAGTCCTCTCTATGGGCAGTCGGTGA | Integrated DNA Technologies (IDT), standard desalting | This study |
| P7_iGuide_PCR2_724 | CAAGCAGAAGACGGCATACGAGATGGTTACGGTTACGTGACTGGAGTCCTCTCTATGGGCAGTCGGTGA | Integrated DNA Technologies (IDT), standard desalting | This study |
| P7_iGuide_PCR2_725 | CAAGCAGAAGACGGCATACGAGATACATCAGGTCACGTGACTGGAGTCCTCTCTATGGGCAGTCGGTGA | Integrated DNA Technologies (IDT), standard desalting | This study |
| P7_iGuide_PCR2_726 | CAAGCAGAAGACGGCATACGAGATGTTGATACGATGGTGACTGGAGTCCTCTCTATGGGCAGTCGGTGA | Integrated DNA Technologies (IDT), standard desalting | This study |
| P7_iGuide_PCR2_727 | CAAGCAGAAGACGGCATACGAGATCAGACACTTCCGGTGACTGGAGTCCTCTCTATGGGCAGTCGGTGA | Integrated DNA Technologies (IDT), standard desalting | This study |
| P7_iGuide_PCR2_728 | CAAGCAGAAGACGGCATACGAGATTCACCATCCGAGGTGACTGGAGTCCTCTCTATGGGCAGTCGGTGA | Integrated DNA Technologies (IDT), standard desalting | This study |
| P7_iGuide_PCR2_729 | CAAGCAGAAGACGGCATACGAGATACCCACCACTAGGTGACTGGAGTCCTCTCTATGGGCAGTCGGTGA | Integrated DNA Technologies (IDT), standard desalting | This study |
| P7_iGuide_PCR2_730 | CAAGCAGAAGACGGCATACGAGATCAGAAGGTGTGGGTGACTGGAGTCCTCTCTATGGGCAGTCGGTGA | Integrated DNA Technologies (IDT), standard desalting | This study |
| P7_iGuide_PCR2_731 | CAAGCAGAAGACGGCATACGAGATGAAGCTTGAATCGTGACTGGAGTCCTCTCTATGGGCAGTCGGTGA | Integrated DNA Technologies (IDT), standard desalting | This study |
| P7_iGuide_PCR2_732 | CAAGCAGAAGACGGCATACGAGATACTAGGATCAGTGTGACTGGAGTCCTCTCTATGGGCAGTCGGTGA | Integrated DNA Technologies (IDT), standard desalting | This study |
| P7_iGuide_PCR2_733 | CAAGCAGAAGACGGCATACGAGATGCTCCTTAGAAGGTGACTGGAGTCCTCTCTATGGGCAGTCGGTGA | Integrated DNA Technologies (IDT), standard desalting | This study |
| P7_iGuide_PCR2_734 | CAAGCAGAAGACGGCATACGAGATTCCCATTCCCATGTGACTGGAGTCCTCTCTATGGGCAGTCGGTGA | Integrated DNA Technologies (IDT), standard desalting | This study |
| P7_iGuide_PCR2_735 | CAAGCAGAAGACGGCATACGAGATTGGCGTCATTCGGTGACTGGAGTCCTCTCTATGGGCAGTCGGTGA | Integrated DNA Technologies (IDT), standard desalting | This study |
| P7_iGuide_PCR2_736 | CAAGCAGAAGACGGCATACGAGATAATCCTCGGAGTGTGACTGGAGTCCTCTCTATGGGCAGTCGGTGA | Integrated DNA Technologies (IDT), standard desalting | This study |
| P7_iGuide_PCR2_737 | CAAGCAGAAGACGGCATACGAGATCTGGACGCATTAGTGACTGGAGTCCTCTCTATGGGCAGTCGGTGA | Integrated DNA Technologies (IDT), standard desalting | This study |
| P7_iGuide_PCR2_738 | CAAGCAGAAGACGGCATACGAGATACCGATTAGGTAGTGACTGGAGTCCTCTCTATGGGCAGTCGGTGA | Integrated DNA Technologies (IDT), standard desalting | This study |
| P7_iGuide_PCR2_739 | CAAGCAGAAGACGGCATACGAGATATGTGCTGCTCGGTGACTGGAGTCCTCTCTATGGGCAGTCGGTGA | Integrated DNA Technologies (IDT), standard desalting | This study |
| P7_iGuide_PCR2_740 | CAAGCAGAAGACGGCATACGAGATTACGTACGAAACGTGACTGGAGTCCTCTCTATGGGCAGTCGGTGA | Integrated DNA Technologies (IDT), standard desalting | This study |
| P7_iGuide_PCR2_741 | CAAGCAGAAGACGGCATACGAGATATCACATTCTCCGTGACTGGAGTCCTCTCTATGGGCAGTCGGTGA | Integrated DNA Technologies (IDT), standard desalting | This study |
| P7_iGuide_PCR2_742 | CAAGCAGAAGACGGCATACGAGATAGCCTGGTACCTGTGACTGGAGTCCTCTCTATGGGCAGTCGGTGA | Integrated DNA Technologies (IDT), standard desalting | This study |
| P7_iGuide_PCR2_743 | CAAGCAGAAGACGGCATACGAGATGCTAAAGTCGTAGTGACTGGAGTCCTCTCTATGGGCAGTCGGTGA | Integrated DNA Technologies (IDT), standard desalting | This study |
| P7_iGuide_PCR2_744 | CAAGCAGAAGACGGCATACGAGATTCTCAGCGCGTAGTGACTGGAGTCCTCTCTATGGGCAGTCGGTGA | Integrated DNA Technologies (IDT), standard desalting | This study |
| P7_iGuide_PCR2_745 | CAAGCAGAAGACGGCATACGAGATGACCCTAGACCTGTGACTGGAGTCCTCTCTATGGGCAGTCGGTGA | Integrated DNA Technologies (IDT), standard desalting | This study |
| P7_iGuide_PCR2_746 | CAAGCAGAAGACGGCATACGAGATTATTCAGCGGACGTGACTGGAGTCCTCTCTATGGGCAGTCGGTGA | Integrated DNA Technologies (IDT), standard desalting | This study |
| P7_iGuide_PCR2_747 | CAAGCAGAAGACGGCATACGAGATGTTCCGGATTAGGTGACTGGAGTCCTCTCTATGGGCAGTCGGTGA | Integrated DNA Technologies (IDT), standard desalting | This study |
| P7_iGuide_PCR2_748 | CAAGCAGAAGACGGCATACGAGATGCGTGTAATTAGGTGACTGGAGTCCTCTCTATGGGCAGTCGGTGA | Integrated DNA Technologies (IDT), standard desalting | This study |
| P7_iGuide_PCR2_749 | CAAGCAGAAGACGGCATACGAGATCTGTAGCTTGGCGTGACTGGAGTCCTCTCTATGGGCAGTCGGTGA | Integrated DNA Technologies (IDT), standard desalting | This study |
| P7_iGuide_PCR2_750 | CAAGCAGAAGACGGCATACGAGATATGCCTCGTAAGGTGACTGGAGTCCTCTCTATGGGCAGTCGGTGA | Integrated DNA Technologies (IDT), standard desalting | This study |
| P7_iGuide_PCR2_751 | CAAGCAGAAGACGGCATACGAGATACCTATGGTGAAGTGACTGGAGTCCTCTCTATGGGCAGTCGGTGA | Integrated DNA Technologies (IDT), standard desalting | This study |
| P7_iGuide_PCR2_752 | CAAGCAGAAGACGGCATACGAGATCTGTTACAGCGAGTGACTGGAGTCCTCTCTATGGGCAGTCGGTGA | Integrated DNA Technologies (IDT), standard desalting | This study |
| P7_iGuide_PCR2_753 | CAAGCAGAAGACGGCATACGAGATCAGTCAGGCCTTGTGACTGGAGTCCTCTCTATGGGCAGTCGGTGA | Integrated DNA Technologies (IDT), standard desalting | This study |
| P7_iGuide_PCR2_754 | CAAGCAGAAGACGGCATACGAGATACTGAGCTGCATGTGACTGGAGTCCTCTCTATGGGCAGTCGGTGA | Integrated DNA Technologies (IDT), standard desalting | This study |
| P7_iGuide_PCR2_755 | CAAGCAGAAGACGGCATACGAGATACGAAGTCTACCGTGACTGGAGTCCTCTCTATGGGCAGTCGGTGA | Integrated DNA Technologies (IDT), standard desalting | This study |
| P7_iGuide_PCR2_756 | CAAGCAGAAGACGGCATACGAGATACCGTCTTTCTCGTGACTGGAGTCCTCTCTATGGGCAGTCGGTGA | Integrated DNA Technologies (IDT), standard desalting | This study |
| P7_iGuide_PCR2_757 | CAAGCAGAAGACGGCATACGAGATAGTCTGTCTGCGGTGACTGGAGTCCTCTCTATGGGCAGTCGGTGA | Integrated DNA Technologies (IDT), standard desalting | This study |
| P7_iGuide_PCR2_758 | CAAGCAGAAGACGGCATACGAGATCCGCACTCAAGTGTGACTGGAGTCCTCTCTATGGGCAGTCGGTGA | Integrated DNA Technologies (IDT), standard desalting | This study |
| P7_iGuide_PCR2_759 | CAAGCAGAAGACGGCATACGAGATTGTGGAAACTCCGTGACTGGAGTCCTCTCTATGGGCAGTCGGTGA | Integrated DNA Technologies (IDT), standard desalting | This study |
| P7_iGuide_PCR2_760 | CAAGCAGAAGACGGCATACGAGATTTAGGCAGGTTCGTGACTGGAGTCCTCTCTATGGGCAGTCGGTGA | Integrated DNA Technologies (IDT), standard desalting | This study |
| P7_iGuide_PCR2_761 | CAAGCAGAAGACGGCATACGAGATTAAGACTACTGGGTGACTGGAGTCCTCTCTATGGGCAGTCGGTGA | Integrated DNA Technologies (IDT), standard desalting | This study |
| P7_iGuide_PCR2_762 | CAAGCAGAAGACGGCATACGAGATCGCGAAGTTTCAGTGACTGGAGTCCTCTCTATGGGCAGTCGGTGA | Integrated DNA Technologies (IDT), standard desalting | This study |
| P7_iGuide_PCR2_763 | CAAGCAGAAGACGGCATACGAGATCGATACACTGCCGTGACTGGAGTCCTCTCTATGGGCAGTCGGTGA | Integrated DNA Technologies (IDT), standard desalting | This study |
| P7_iGuide_PCR2_764 | CAAGCAGAAGACGGCATACGAGATTTGAAATCCCGGGTGACTGGAGTCCTCTCTATGGGCAGTCGGTGA | Integrated DNA Technologies (IDT), standard desalting | This study |
| **MiSeq Sequencing Primers** |  |  |  |
| Index1_Seq_Primer | ATCACCGACTGCCCATAGAGAGGACTCCAGTCAC | Integrated DNA Technologies (IDT), standard desalting | PMID: 25513782 |
| Read2_Seq_Primer | GTGACTGGAGTCCTCTCTATGGGCAGTCGGTGAT | Integrated DNA Technologies (IDT), standard desalting | PMID: 25513782 |
