## Supplementary Table 12 for "An integrated enzymatic and computational pipeline for quantifying off-target base-editing"

**Supplementary Table 12. Key reagents**

| **Reagent** | **Company** | **Cat.no.** |
| --- | --- | --- |
| CD4 CliniMACS beads | Miltenyi Biotec | 200-070-213-276-01 |
| CD8 CliniMACS beads | Miltenyi Biotec | 200-070-215-275-01 |
| CD34 microbeads | Miltenyi Biotec | 130-046-703 |
| Supplemented CTS OpTmizer T Cell Expansion Media | Gibco | A10484-02 |
| X-Vivo | TheraPeak Lonza | BP04-744Q |
| Human Ab Serum plasma-derived male pooled | Bioivt | HUMANABSRMP-HI-1 |
| Glutamax | Gibco | 35050-061 |
| Penicillin Streptomycin | Gibco | 15140-122 |
| IL-7 | Peprotech | AF-200-07-100UG |
| IL-15 | Peprotech | AF-200-15-100UG |
| StemSpan SFEM II | StemCell Technologies | 9655 |
| hSCF | Peprotech | AF-300-07-100UG |
| FLT3L | Peprotech | AF-300-19-100UG |
| TPO | Peprotech | AF-300-18-100UG |
| IL6 | Peprotech | AF-200-06-100UG |
| UM729 | StemCell Technologies | 72332 |
| Dynabeads Human T-Activator CD3/CD28 | Gibco | 11132D |
| P3 Primary Cell Nucleofector Solution with Supplement 1 | Lonza | PBP3-02250 |
| Hyclone buffer | Cytiva | EPB1 |
| Quick-DNA Miniprep Plus Kit | Zymo Research | D4069 |
| Phusion DNA Polymerase | New England Biolabs | M0530L |
| DNA Clean & Concentrator-5 kit | Zymo Research | D4013 |
| rhAmpSeq CRISPR Library Kit | IDT | 10007318 |
| ABE8e-SpCas9 (D10A)-NGG mRNA, clean-cap AG, N1-methyl-pseudouridinylated | TriLink | L-7010 |
| ABE8e-SpCas9^WT^-NGG mRNA, clean-cap AG, N1-methyl-pseudouridinylated | TriLink | L-7010 |
| Clean-cap Cas9 mRNA | TriLink | L-7606 |
