## Supplementary Table 1 for "An integrated enzymatic and computational pipeline for quantifying off-target base-editing"

**Supplementary Table 1. Examples of published methods for monitoring off-target base editing**

| **In-silico assay** | **Method Description** | **Method Readout** | **Reference** |
| --- | --- | --- | --- |
| Cas-OFFinder | Cut site nomination via custom pattern searching algorithm. | Table with positions including mismatches and bulges. | PMID: 24463181 |
| CCTOP | Cut site nomination via alignment with Bowtie. | Table with positions, PAM, mismatches, and genomic region annotation. | PMID: 25909470 |
| CHOPCHOP | Cut site nomination by alignment with Bowtie. | Table with positions and number of mismatches. | PMID: 31106371 |
| BEdeepoff | Cut site nomination by alignment followed by cut efficiency prediction using a recurrent neural network. | Table with positions and number of mismatches, bugles, and predicted cut efficiency. | PMID: 37660097 |
| COSMID | Cut site nomination using FetchGWI. Scoring based on mismatches. | Table with positions, mismatches, and score. | PMID: 25462530 |
| CRISPOR | Cut site nomination via BWA. Scoring based on gap and mismatch penalties. | Table with positions, two off-target scoring methods, and genomic region annotation. | PMID: 29762716 |
| CRISPRme | Cut site nomination via custom alignment algorithm. Scores via systems from CRISTA and cutting frequency determination. | Table with positions, PAM, mismatches, bulges, two scoring method, variant detection, and genomic region annotation | PMID: 36522432 |
| CRISPRoff | Cut site scoring using free energy changes. | Table with position, score, gene annotation. | PMID: 36271848 |
| CRISTA | Cut site nomination using BWA. Random-forest based method for predicting cut effieincy. | Table with position, score, PAM, bulges, mismatches, and physical characteristics. | PMID: **29036168** |
| DeepBE | Base edit efficiency prediction using a convolutional neural network. | Table with predicted edit efficiency and inferred proportion of edits at the protospacer level. | PMID: **37188916** |
| BEDICT | Base edit efficiency prediction using a transformer-based autoregressive sequence-to-sequence model. | Table with per-base predicted edit efficiencies. | **PMID: 34433819** |
| **Biochemical assay** |  |  |  |
| nDigenome-Seq | WGS of gDNA that has been treated with ABE:gRNA RNP complexes. | OTS are identified as reads with staggered and straight alignments. | PMID: 32941652 |
| EndoV-Seq | WGS of gDNA that has been first treated with ABE:gRNA RNP complexes, followed by EndoV treatment, which captures the inosine intermediate, which results in DSBs. | OTS are identified as reads with staggered and straight alignments. | PMID: 30622278 |
| CHANGE-Seq-BE | gDNA is tagmented using transposase and circularized, followed by incubation with ABE:gRNA RNP complexes, and EndoV. Which generates linearized sequences if digest is successful. | OTS are identified as they lead to linearized gDNA circles that are used for library prep and high-throughput sequencing. | PMID: 38585919 |
| Selict-Seq | gDNA treatment with ABE:gRNA RNP complexes, capturing of dI with biotinylated EndoV after crosslinking? and pulldown. | OTS are identified by high-throughput sequencing of enriched fragments. |  |
| Tracking-Seq | Editing in cells and gRNA isolation or gDNA treatment with ABE:gRNA RNP complexes, followed by RPA2 incubation that binds and protects ssDNA. dsDNA and protein digestion to enrich ssDNA. | OTS are identified by high-throughput sequencing of enriched fragments. | PMID: 38956324 |
| TOPO-Seq | gDNA is tagmented using transposase and circularized (like CHANGE-Seq /BE), followed by incubation with Cas9:gRNA or ABE8e:gRNA RNP complexes and EndoV under a gradient of ethidium bromide, which supercoils DNA, and generates linearized sequences if the digest is successful. | OTS are identified as they lead to linearized gDNA circles that are used for library prep and high-throughput sequencing. | PMID: 40175512 |
| **Cellular assay** |  |  |  |
| (BE)-iGUIDE-Seq | Editing was performed in cells using a DSB active ABE or CBE while also introducing dsODN. gDNA isolation, fragmentation, and enrichment of dsODN-tagged genomic regions. | OTS are identified by high-throughput sequencing of enriched fragments and base editing in the vicinity. | iGUIDE-Seq PMID: 30654827 |
| Detect-Seq | Editing in cells and gDNA was isolated and fragmented. dU (intermediate generated by CBEs) was labeled with biotin for pulldown and enrichment. | OTS are identified by high-throughput sequencing of enriched fragments | PMID: 34099937 |
| Induce-Seq | Editing in cells and in situ labelling ds/ssDNA with an adapter. Fragmentation and addition of sequencing adapters, followed by enrichment on the flow cell. | OTS are identified by high-throughput sequencing of enriched fragments | PMID: 35810156 |
